# Structure and dynamic association of an assembly platform subcomplex of the bacterial type II secretion system

**DOI:** 10.1101/2022.07.16.500195

**Authors:** Régine Dazzoni, Yuanyuan Li, Aracelys López-Castilla, Sébastien Brier, Ariel Mechaly, Florence Cordier, Ahmed Haouz, Michael Nilges, Olivera Francetic, Benjamin Bardiaux, Nadia Izadi-Pruneyre

## Abstract

Type II secretion systems (T2SS) allow diderm bacteria to secrete hydrolytic enzymes, adhesins or toxins important for growth and virulence. In T2SS, secretion of folded proteins from the periplasm to the cell surface requires assembly of periplasmic filaments called pseudopili. Like the related type IV pili, pseudopili are polymerized in the inner membrane through addition of subunits at the filament base, mediated by the essential assembly platform (AP). To understand the structure and molecular role of the AP, we focused on its components PulL and PulM from the *Klebsiella oxytoca* T2SS. By combining biophysical methods, NMR and X-ray crystallography we studied the structure and associations of their periplasmic domains. We describe the first structure of the heterodimer complex formed by the PulL and PulM ferredoxin-like domains and show how their structural complementarity and plasticity favor their association during the secretion process. Cysteine scanning and cross-linking of transmembrane segments provided additional constraints to build a structural model of the PulL–PulM complex and assembly in the cellular context. Together with the relative abundance of PulL, PulM and their partners our findings suggest a model of the AP as a dynamic hub that orchestrates pseudopilus polymerization.

## Introduction

Gram-negative bacteria have developed multiple protein secretion systems that are important for their survival and pathogenesis (Maffei *et al*., 2017). Among these, the type II secretion system (T2SS), discovered in the late 1980s (d’Enfert *et al*., 1987), is one of the most widespread and relevant from biomedical and environmental standpoints. It allows the bacterium to secrete fully folded proteins with a wide range of functions – toxins, adhesins, cytochromes and hydrolytic enzymes (Cianciotto and White, 2017).

The T2SS is a transmembrane nano-machine composed of twelve to fourteen proteins designated here using the Gsp (General secretory pathway) nomenclature (Pugsley, 1993). It is organized into four sub-complexes (reviewed in (Naskar *et al*., 2021)): (1) an outer membrane secretin channel (GspD); (2) a periplasmic pseudopilus filament composed of a non-covalent polymer of major pseudopilin GspG and minor pseudopilins GspH, I, J, K; (3) an assembly platform (AP) composed of GspL, M, C and F (Py *et al*., 2001) ; and (4) a cytosolic ATPase GspE. This system has a common evolutionary origin (Hobbs and Mattick, 1993, Peabody *et al*., 2003, Denise *et al*., 2019) and shares a similar architecture with type IV pili (T4P) and archaeal flagella and pili, together forming the superfamily of type IV filament assembly systems (Berry and Pelicic, 2015).

Although the general architecture of the T2SS has been extensively investigated, the biogenesis of this nanomachine and the secretion mechanism are still largely unknown. It is assumed that the substrate secretion is coupled with the polymerization of the pseudopilus in the periplasm, which is driven by the assembly platform (AP) complex. However, how pseudopili are assembled and how they drive secretion remains elusive. Information on the interaction mode of AP components and their structure is essential to understand these mechanisms.

Here we focused on two essential AP components GspL and GspM from *Klebsiella oxytoca* T2SS called respectively PulL and PulM, with reference to the lipoprotein pullulanase (PulA), the only identified exoprotein secreted by this system. This enzyme degrades branched maltotriose polymers, allowing them to be taken up and used as nutrient (d’Enfert *et al*., 1987). GspL and GspM have similar domain organization suggesting a common evolutionary origin. Both are predicted to insert in the inner membrane *via* a single hydrophobic helix that is followed by a periplasmic α-helical region and a C-terminal globular domain. X-ray crystallography has provided structural information for the periplasmic domains of GspM (Abendroth *et al*., 2005) and GspL (Abendroth *et al*., 2009, Fulara *et al*., 2018). In addition, GspL has an N-terminal cytoplasmic domain, which forms a stable complex with the ATPase GspE, as demonstrated in the *Vibrio cholerae* T2SS (Abendroth *et al*., 2005). This interaction anchors the GspE hexamer to the cytoplasmic base of the AP complex. *In vivo*, GspM forms a complex with GspL, protecting it from degradation, as shown in *V. cholerae* (Sandkvist *et al*., 1999). Deleting *pulM* gene in *K. oxytoca* also leads to degradation of PulL, which is in turn required for the stability of the ATPase PulE (Possot *et al*., 2000). An electron-microscopy study of a purified T2SS subcomplex from *K. pneumoniae* suggested a C6 symmetry of the GspE-GspL-GspM complex (Chernyatina and Low, 2019). However, neither that study, nor the *in situ* analysis of the *Legionella pneumophila* T2SS by cryo-tomography (Ghosal *et al*., 2019) were able to provide a clear view of the L and M complex architecture.

Here we studied the structure and the assembly of PulL and PulM by focusing first on their periplasmic globular C-terminal domains (CTD). Due to their dynamics, their structural study was only possible by an integrative approach. By combining native mass spectrometry, NMR and X-ray crystallography we showed that while each protein alone is a homodimer in solution, it forms a heterodimer in the presence of its partner. Interestingly, the presence of the partner drives the exchange of homodimer interfaces toward the formation of the heterodimer, showing their structural complementarity and plasticity. To our knowledge, this is the first reported high-resolution structure of a heterodimer complex of the AP components. To reach a comprehensive view of PulL-PulM assembly in the cellular and membrane context, we used bacterial two-hybrid (BACTH), cysteine crosslinking and *in vivo* functional assays. Together our structural and functional data allow us to propose a model of the dynamic association of AP proteins and of the way it drives the pseudopilus formation during the protein secretion processes.

## Results

Previous studies report evidence for a direct interaction between the AP components PulL and PulM in *K. oxytoca* (Possot *et al*., 2000, Nivaskumar *et al*., 2016). In the *Dickeya dadantii* T2SS, bacterial two-hybrid and cysteine crosslinking assays suggested the formation of a complex between GspL and GspM periplasmic C-terminal domains (Lallemand *et al*., 2013). However, their assembly mode is still not understood at the molecular and atomic levels. Here, to study their structure and interaction, we produced and purified the predicted globular C-terminal domains of PulL (PulL_CTD_, residues 312-398) and PulM (PulM_CTD_, residues 79-161).

### PulL_CTD_ structure

First by NMR and analytical ultracentrifugation (AUC), we determined the oligomerization state of PulL_CTD_ at concentrations ranging from 10 to 300 μM (Figure S1 and Supplementary Material). PulL_CTD_ exists in monomer-dimer equilibrium depending on the protein concentration. At the concentration used for NMR experiments (300-400 μM), it is mainly in dimeric form. Its ^1^H-^15^N HSQC spectrum was used for an initial structural analysis. In this spectrum, each signal arises from the backbone amide group of one residue. The number of observed signals in the ^1^H-^15^N HSQC spectrum of PulL_CTD_ (82) is consistent with the number of its backbone amide groups (83), and thus with a symmetrical homodimer (Figure 1A). To solve the structure of the PulL_CTD_ dimeric form, we first determined the structure of its monomer by NMR. An ensemble of 15 monomeric structures was calculated based on chemical shift assignments obtained previously (Dazzoni *et al*., 2021) and NOESY experiments by using the program ARIA (Allain *et al*., 2020) as detailed in Materials and Methods. The ensemble of PulL_CTD_ monomer structures (Figure 1B) was calculated with 996 restraints, and presents a backbone RMSD of 0.6 Å for its ordered part (Table S1). PulL_CTD_ displays a ferredoxin-like fold with 2 helices, α1 (from S319 to D332) and α2 (from S359 to R372), and an anti-parallel β-sheet formed by 4 β-strands, β1 (from I336 to D344), β2 (from N348 to A356), β3 (from F373 to Q376) and β4 (from I387 to G395) (Figure 1B).

**Figure 1.**
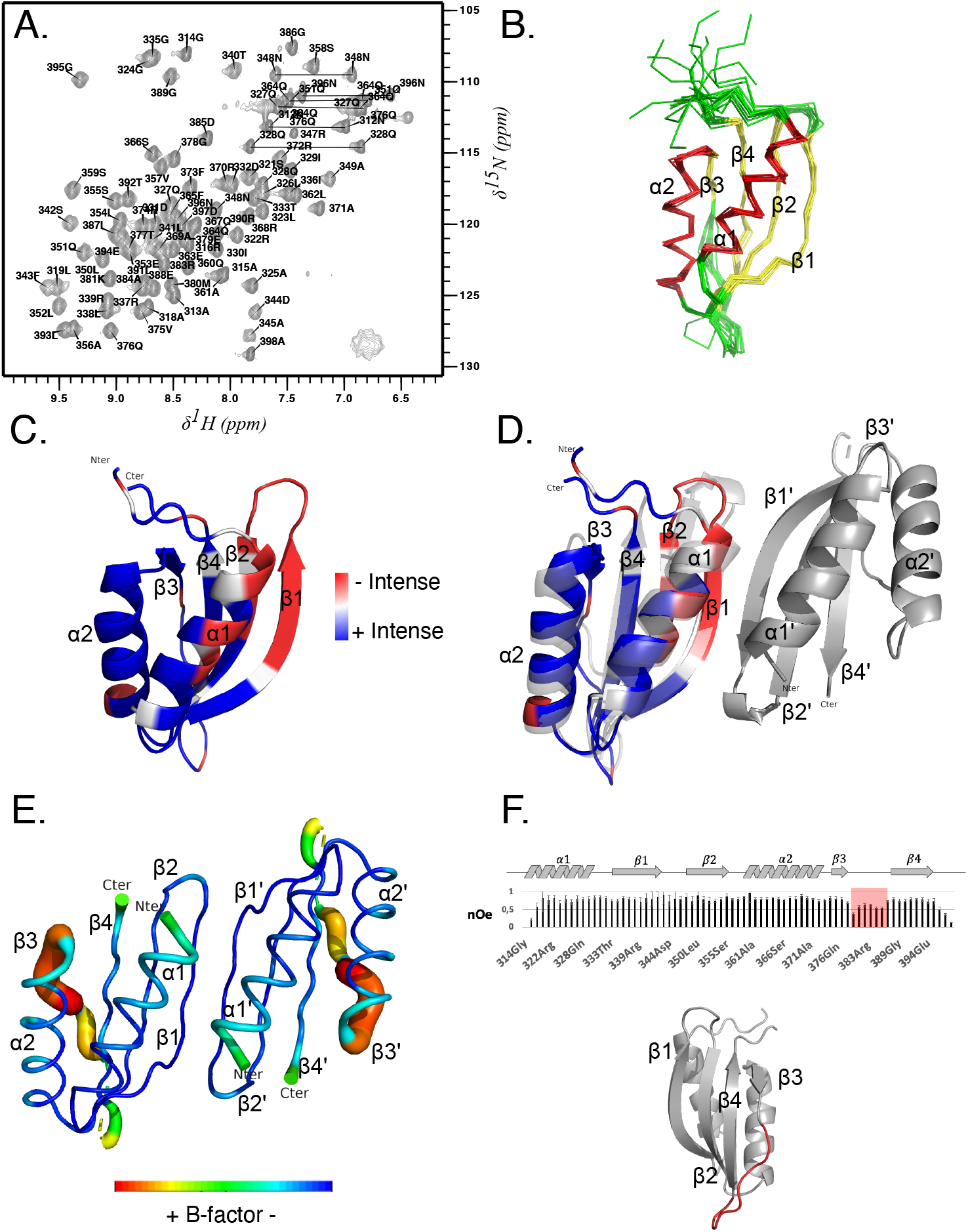
PulL_CTD_ homodimer structure. **A**. ^1^H-^15^N HSQC NMR spectrum of PulL_CTD_ (400 μM in 50 mM HEPES buffer pH 6.5, 50 mM NaCl). Backbone resonance assignments are indicated in one-letter amino acid code and side chain NH_2_ peaks of Asn (N) and Gln (Q) are connected by horizontal lines. **B**. NMR structure ensemble of PulL_CTD_ exhibiting a ferredoxin-like fold (α1-β1-β2-α2-β3-β4; α-helix in red, β-sheet in yellow, loops and turns in green). **C**. Cartoon representation of the lowest-energy NMR structure of the PulL_CTD_ monomer. The intensity of the ^1^H-^15^N HSQC signals of PulL_CTD_ are reported with a specific colour code on the structure from blue to red for decaying peak intensity, outlining the likely homodimeric interface. **D**. Superposition of the PulL_CTD_ NMR structure (coloured as in C) and the dimer structure of PulL_CTD_ obtained by X-ray crystallography in grey (PDB ID: 8A9W). The twofold symmetry axis is shown as a black ellipse. **E**. Representation of the B-factor per residues of PulL_CTD_ in the X-ray crystallography dimer structure, from blue to red for increasing B-factor. **F**. ^1^H–^15^N heteronuclear NOE values along the sequence of PulL_CTD_. The red box indicates the lower values corresponding to the most dynamic region (ps-ns time scale), located on the β3-β4 loop shown in red in the cartoon model.

To determine the structure of the dimeric form, ^13^C/^15^N edited-filtered NOESY experiments were performed to collect intermolecular distance restraints between two protomers of PulL_CTD_. Unfortunately, no intermolecular cross-peaks could be observed in these experiments. The absence of signal in such spectra can be due to conformational exchange of the corresponding residues at the μs-ms NMR time scale. Dynamics at this timescale has been already suspected in the ^1^H-^15^N HSQC spectrum for several residues displaying lower peak intensity leading to weak or missing peaks in 3D NMR experiments (Dazzoni *et al*., 2021). When mapping the NMR signal intensity on the PulL_CTD_ monomer structure (Figure 1C), it appears that these residues are mostly located on the same face of the molecule (α1, β1 and β2) suggesting conformational exchange of these residues between monomeric and dimeric forms.

Without any inter-protomer distance restraints, it was not possible to determine the structure of the dimeric form by NMR; we thus used X-ray crystallography and solved the structure of PulL_CTD_ at 1.8 Å resolution. The summary of data collection and refinement statistics is shown in Table S2. The structure of PulL_CTD_ exhibits a nearly identical ferredoxin-like fold (α1-β1-β2-α2-β3-β4) as observed in the NMR structure (Figure 1D). However, the β3 strand (residues F373-Q376) is not fully formed in the X-ray structure. As shown in Figure 1E, higher B-factor values are observed in this region compared to the rest of the protein, reflecting a disordered β-strand. Consistently, higher flexibility on the ps-ns time scale in this strand is evidenced by the lower ^1^H–^15^N heteronuclear NOE values (Figure 1F).

Only one possible dimer interface with a C2 symmetry was found, across two asymmetric units in the crystal lattice and with a buried surface area of 526 Å^2^. This dimeric interface occurs between the α1 helix and the β1 strand oriented in an anti-parallel fashion (Figure 1D). Although the interaction surface is quite large relative to the size of the domain, only 6 hydrogen bonds are observed at the dimerization interface: one between A345 and R339 of β1 and β1’; and two between F343 and L341 in both protomers (see also Figure 6G below). There are also a few interactions involving α1 helices, notably hydrophobic contacts between the I320 side-chains of each protomer. This small number of interprotein contacts explains the short-lived assemblies (Janin *et al*., 2007) and the monomer/dimer exchange of PulL_CTD_ in solution observed by NMR and AUC.

### PulM_CTD_ structure

PulM_CTD_ behaves as a stable dimer in solution at low and high concentrations (Figure S1 and Supplementary Material). To determine its structure, we performed NMR experiments and crystallization trials in parallel. We obtained crystals in two days, with an asymmetric unit containing seven virtually identical PulM_CTDs_ (average RMSD 0.28 Å for Cα atoms). The final structure was refined at 1.5 Å of resolution (Figure 2A, B). The summary of data collection and refinement statistics is shown in Table S2. The structure of the PulM_CTD_ protomer exhibits a ferredoxin-like fold (α1-β1-β2-α2-β3-β4). The first helix α1 extends from P82 to H93, followed by β1 from V98 to Q103, β2 from R106 to V111, the α2 helix from F116 to A129, β3 from A134 to A140 and β4 from V148 to E156.

**Figure 2:**
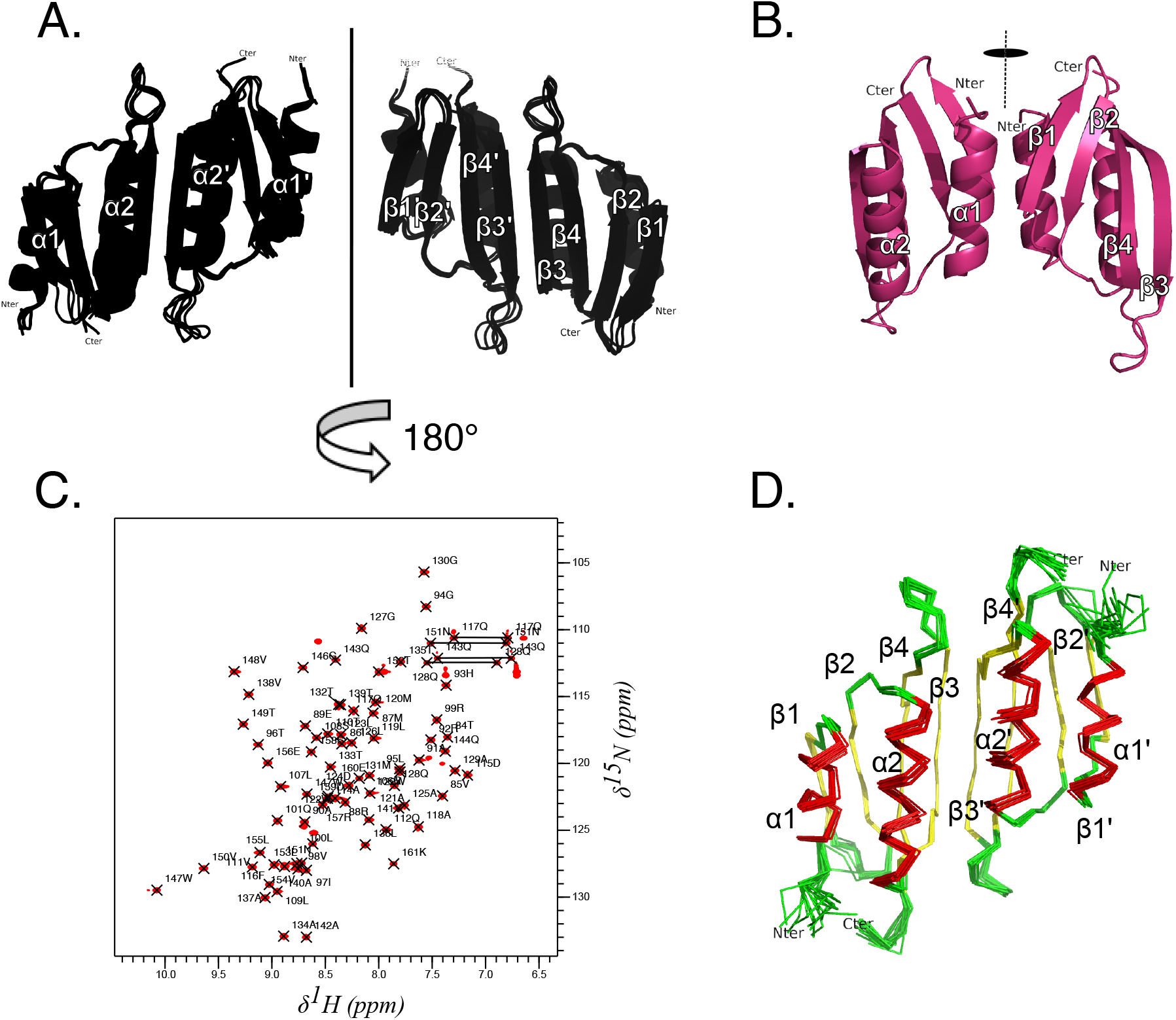
PulM_CTD_ homodimer structure. **A**. Main topology (form A) found in the PulM_CTD_ crystal. The four homodimers observed in the asymmetric unit are superimposed. **B**. Minor topology (form B) found in the PulM_CTD_ crystal, represented by a single homodimer. The vertical lines indicate the twofold symmetry axis. **C**. ^1^H-^15^N HSQC NMR spectrum of PulM_CTD_ in 50 mM HEPES pH 7.0, 50 mM NaCl. Backbone resonance assignments are indicated in one-letter amino acid code. Side chain NH_2_ signals of Asn (N) and Gln (Q) are connected by horizontal lines. **D**. NMR structure ensemble of the PulM_CTD_ homodimer. Helices are coloured in red, β-strands in yellow and turns/loops in green.

Across the crystal symmetry, two dimer topologies of PulM_CTD_ were found, named form A and form B. Form A was represented by four virtually superimposable anti-parallel dimers (average RMSD= 0.7 Å), with an interface involving α2-β3 regions (Figure 2A). Form B was represented by a single parallel homodimer with the α1-β1 interface (Figure 2B).

These alternative dimeric forms A and B have, respectively, a buried surface area of 643.7 Å^2^ and 462.9 Å^2^ across the dimerization interface, as calculated by using the PISA server (Krissinel and Henrick, 2007). The difference in buried interface is small, but significant considering the small size of the domain with a total surface area of 4,500 Å^2^. The A form is thus more favorable from a structural aspect, considering its larger buried interface and the fact that it is also the most abundant form in the crystal (80%).

Crystal packing can favor either form, therefore we used NMR spectroscopy to determine which dimeric form of PulM_CTD_ exists in solution. We first determined the structure of PulM_CTD_ in its monomeric form and then collected intermolecular cross-peaks by using ^13^C- and ^15^N-filtered NOESY experiments to determine the dimer structure. A total of 1623 intramolecular and 25 intermolecular restraints were used to calculate the structure of PulM_CTD_ dimer with a backbone RMSD of 0.69 Å for the 15 lowest energy structures (Figure 2D and Table S3). The PulM_CTD_ monomer structure exhibits a ferredoxin-like fold (α1-β1-β2-α2-β3-β4). The α1 helix comprising residues T84 to H93, is followed by β1 strand from V98 to Q101 and β2 strand from L107 to V111, the second helix α2 from F116 to A129, β3 stand from M131 to A140 and strand β4 from V148 to R157. In the solution structure, the dimer interface occurs between the α2 helix and the β3 strand of the two protomers, oriented in an antiparallel fashion (Figure 2D). This dimer in solution is therefore similar to the form A obtained by X-ray crystallography (RMSD of backbone atoms 1.6 Å between the PulM_CTD_ NMR dimer structure and the crystallographic form A). While the monomers show an almost identical structure (backbone RMSD of 1.1 Å for secondary structure elements), the most noticeable difference between the NMR and X-ray crystallographic structures of the PulM_CTD_ dimer is a slight displacement of the α2 helices, 2.3 Å on average along the helix axis (Figure S2).

Although the dimerization interface and the inter-protomer interactions are similar in both NMR and X-ray structures, the number of hydrogen bonds between the monomers in the β3/β3’ anti-parallel β sheet is different: 7 in the X-ray structure versus 6 in the NMR structure ensemble. Six of them are between the two β3 strands and involve the residues L136-V138 and A140-A134. In the X-ray structure an additional hydrogen bond links the α2 residues Q117 and D124 of each monomer. In addition, a series of hydrophobic contacts between F116, M120, L123, L136, and V138 contribute to the stabilization of the α2-β3 interface as observed in both NMR and X-ray structures (see also Figure 6B below).

In T4P assembly systems, the CTD of the PulM orthologue PilO also forms homodimers but with β-sheets facing the opposite sides in the two protomers (Sampaleanu *et al*., 2009, Leighton *et al*., 2016, 2018).

### PulL_CTD_-PulM_CTD_ interaction

To gain insight into the PulL_CTD_-PulM_CTD_ assembly and mode of interaction, we analysed their interaction by NMR. The ^1^H-^15^N HSQC spectra were recorded for the mixtures of a labelled protein and its unlabelled partner. Spectral variations in the presence of a partner is a strong indicator of an interaction. The titration experiments of each protein with its unlabelled partner up to three-fold molar ratio, indicated a stoichiometry of 1 PulL_CTD_ for 1 PullM_CTD_ (Figure S1B). The rotational correlation time τ _c_ for the PulL_CTD_-PulM_CTD_ complex (between 10.4 and 10.8 ns) was also consistent with a 1:1 stoichiometry as a heterodimer (Figure S1A). For a more accurate characterization of the oligomeric state, we employed native mass spectrometry approach. Both PulL_CTD_ (Figure 3A) and PulM_CTD_ (Figure 3B) are present as homodimers in accordance with their expected molecular weight (expected/measured MW: 19 069.48 Da/19 069.15 Da for PulL_CTD_; 17 861.42 Da/17 861.87 Da for PulM_CTD_). The mass spectrum of an equimolar mixture of PulL_CTD_ and PulM_CTD_ showed the presence of an additional species with a well-resolved charge state series, from +7 (*m/z* 2638.9) to +9 (*m/z* 2052.7) (Figure 3C). The measured molecular weight (18 465.63 Da) fits perfectly with the expected mass of the PulL_CTD_-PulM_CTD_ heterodimer (18 465.95 Da). The presence of both homo- and heterodimeric species in the mixture can be due to molecular exchange between these two conformations at this concentration. No higher oligomeric states were detected, confirming that PulL_CTD_ and PulM_CTD_ form heterodimers under our experimental conditions.

**Figure 3.**
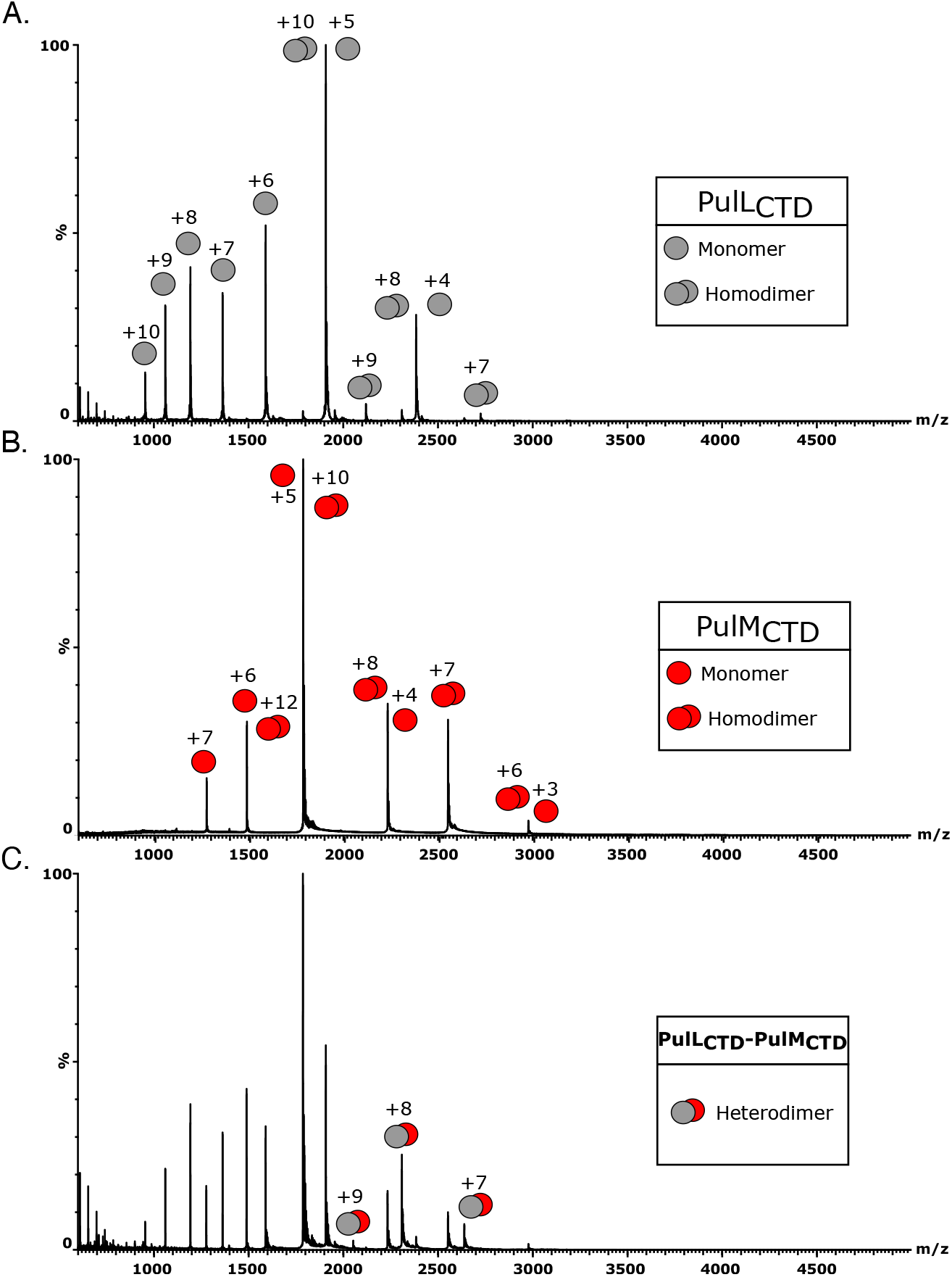
Oligomeric state of the PulL_CTD_-PulM_CTD_ complex. Native mass spectra of PulL_CTD_ (**A**) and PulM_CTD_ (**B**) alone and in an equimolar mixture of PulL_CTD_ and PulM_CTD_ (**C**). From mass spectra we observe m/z corresponding to the presence of monomeric, homodimeric and heterodimeric species (annotated in grey for PulL_CTD_ and in red for PulM_CTD_). Charge states are indicated for all species.

To further understand the PulL_CTD_-PulM_CTD_ interaction mode, we solved the crystal structure of the PulL_CTD_-PulM_CTD_ complex. A summary of the crystallographic parameters and data, as well as refinement statistics, are shown in Table S2. The asymmetric unit contains three PulM_CTD_ and one PulL_CTD_ molecules, organized in an arc-like arrangement, named: 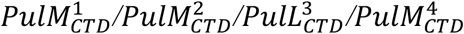 (Figure 4). PISA analysis of molecular interfaces found within the crystal revealed two main interacting surfaces within the asymmetric unit, and a smaller one (interface area < 500 Å) between 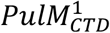 and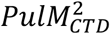. This PulM_CTD_ homodimer corresponds to the minor B form that was also observed in the crystal of PulM_CTD_. Another interaction occurs between two molecules of PulM_CTD_ 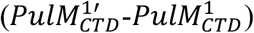 from two neighboring asymmetric units, exhibiting a buried interface of 601.6 Å^2^. This homodimer is equivalent to the one observed by NMR and in the crystallographic form A of PulM_CTD_.

**Figure 4.**
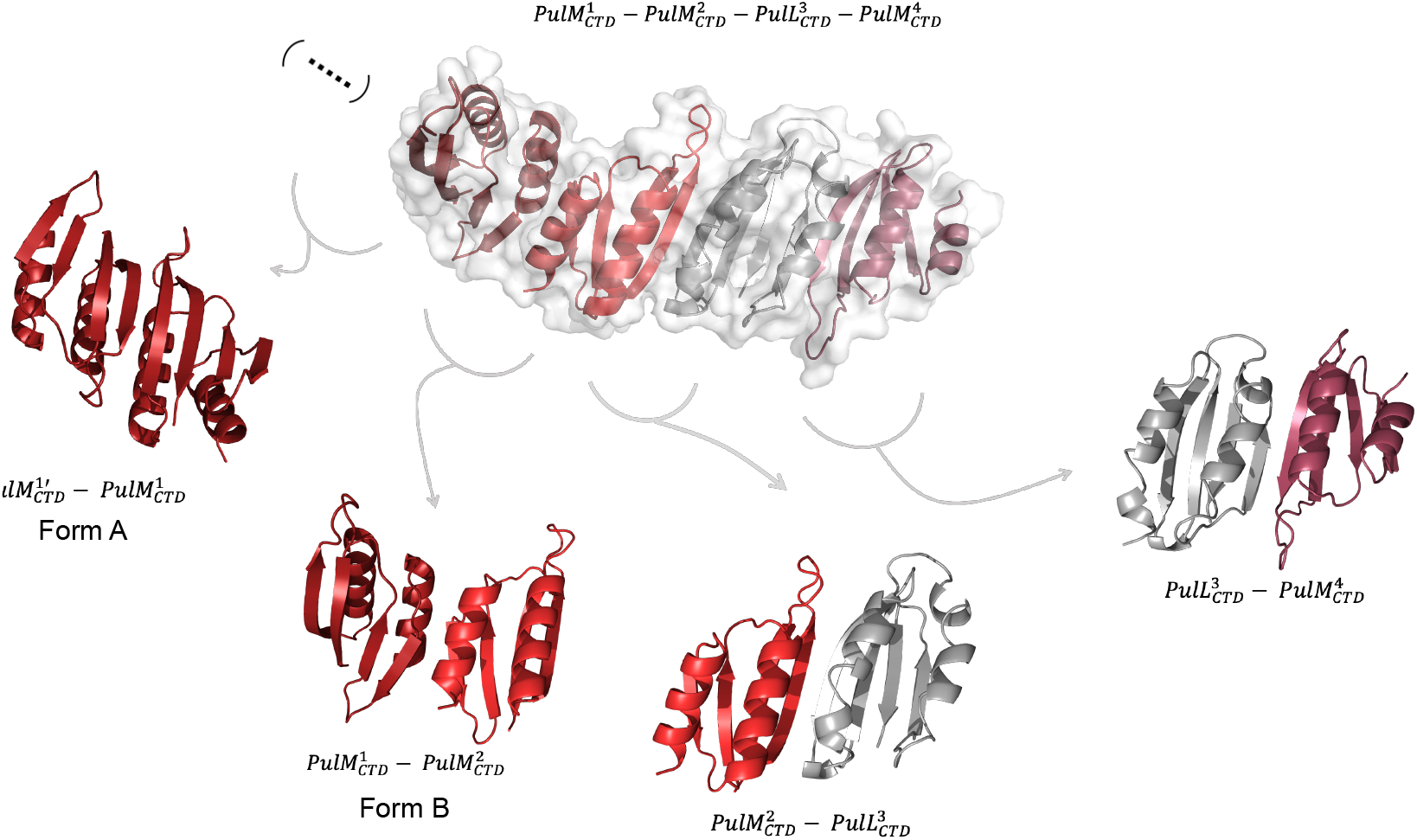
PulL_CTD_-PulM_CTD_ complex structure determined by X-ray crystallography. Top: the asymmetric unit composed of: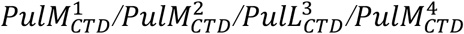. Bottom: Different homo/heterodimeric forms found in the asymmetric unit and crystal lattice. The different PulM_CTD_ subunits are colored in shades of red and PulL_CTD_ in grey.

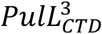 shares two different interfaces with either 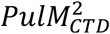 or 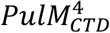 subunits in the asymmetric unit with buried interfaces of 639 Å^2^ and 544 Å^2^, respectively. As the difference is small, we used two additional methods to score their interfaces *i*.*e*. Prodigy-Crystal (Jiménez-García *et al*., 2019) and ClusPro-DC (Yueh *et al*., 2017) which allowed us to conclude that the 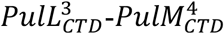 heterodimer is more likely to represent a biologically relevant arrangement (Table S4).

To complete the analysis of PulL_CTD_-PulM_CTD_ interaction in solution, we compared ^1^H-^15^N HSQC spectra of the ^15^N-PulM_CTD_ and ^15^N-PulL_CTD_ each alone and in the presence of its unlabeled partner (Figure 5A, B) and measured the corresponding chemical shift perturbation (CSP) of backbone amide signals. For PulL_CTD_, positions displaying the highest CSPs (>1.5σ above the mean CSP) were residues L319, I320, S321 and L326 of the α1 helix; the β-sheet was also affected, namely the β1-β2 strands (from R337 to A356) and the middle of β4 at the C-terminus (from L391 to G395) (Figure 5C, E). On PulM_CTD_, the highest CSPs (>1.5σ) are observed within the α2 helix (from Q117 to Q128), the β3 strand (from G130 to V138) and the loop connecting β3 to β4 (V141 to V148). As observed in Figure 5E and F, the majority of the highest CSPs for both proteins are located at their respective homodimerization interface. This suggests that the PulL_CTD_-PulM_CTD_ complex has a single binding interface and can form only binary complexes, which is consistent with the mass spectrometry data. This interface of dimerization in solution corresponds to the 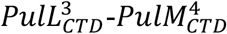 that we identified as the most relevant one in the PulL_CTD_-PulM_CTD_ crystal structure.

**Figure 5.**
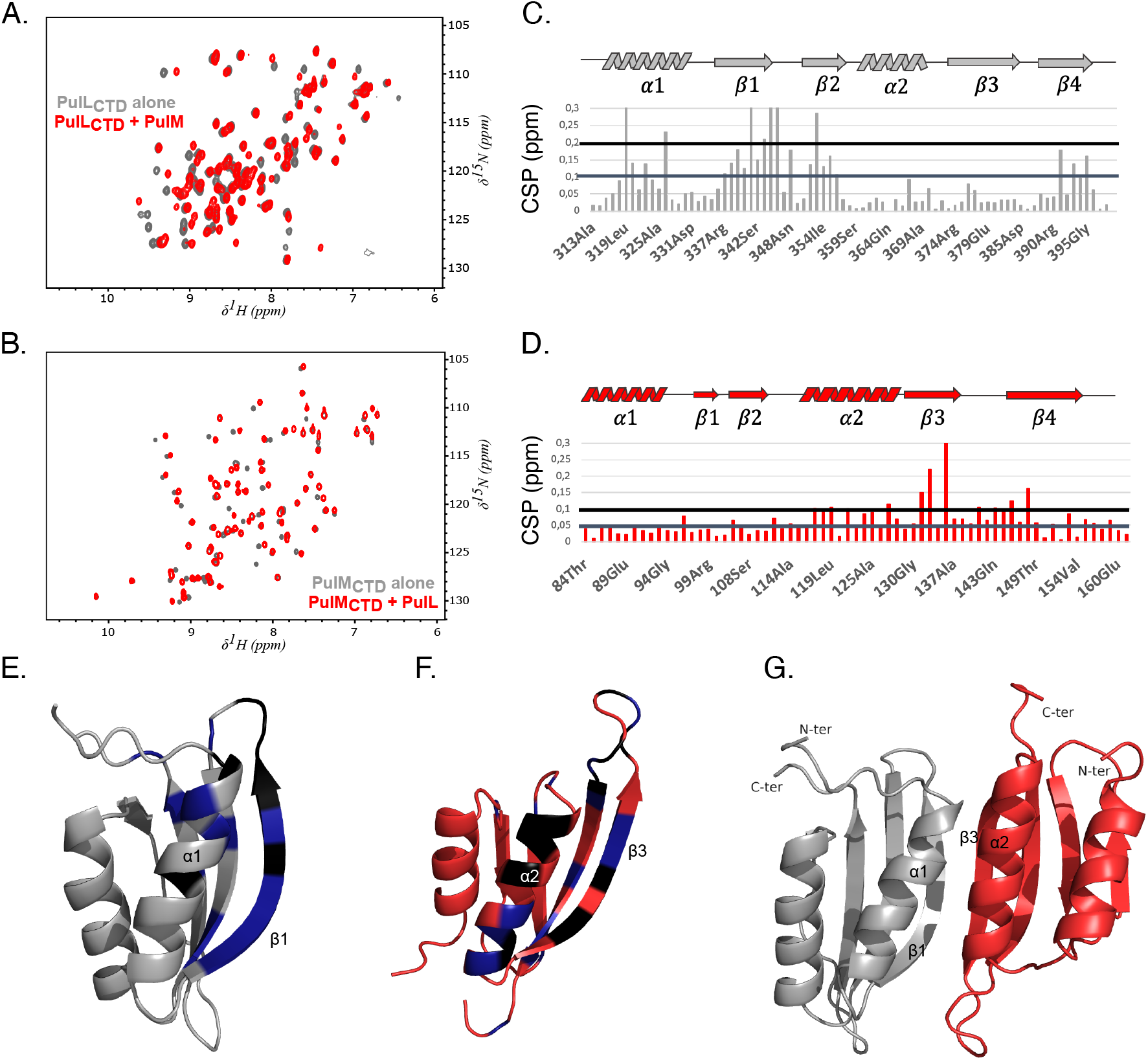
NMR analysis of PulL_CTD_-PulM_CTD_ interface. **A**. Overlay of ^1^H-^15^N HSQC spectra of PulL_CTD_ in the absence (grey) and presence (red) of unlabeled PulM_CTD_. **B**. Overlay of ^1^H-^15^N HSQC spectra of PulM_CTD_ in absence (grey) and presence (red) of unlabeled PulL_CTD_. **C**. Chemical shift perturbation (CSP) of PulL_CTD_ backbone amide signals as a function of residue number. The grey lines indicate CSP value higher than 3 or 1.5 standard deviations (α) from the mean CSP. **D**. CSP of PulM_CTD_ backbone amide signals as a function of residue numbers. The grey lines indicate CSP value higher than 3α or 1.5α from the mean CSP. **E**. CSP values reported on the NMR structure of PulL_CTD_. In black, CSP higher than 3α; in blue CSP between 1.5 and 3α. **F**. CSP values reported on the NMR structure of PulM_CTD_. Same color code as in E. **G**. Model of the PulL_CTD_-PulM_CTD_ complex generated with the docking program HADDOCK and using CSP as restraints.

We could not determine the structure of PulL_CTD_-PulM_CTD_ complex by NMR, since no signal corresponding to intermolecular NOE could be observed, although various conditions were tested (temperature, magnetic fields and pulse sequences). The absence of intermolecular NOEs is most likely due to the dynamics of the heterodimer interface at the μs-ms time scale. Therefore, we used the CSPs to guide modelling of the complex with the HADDOCK docking program (van Zundert *et al*., 2016, Honorato *et al*., 2021). The CSP analysis was complemented by residue co-evolution predicted by EVcoupling (Hopf *et al*., 2014) highlighting two compatible pairs of co-evolving residues across the PulL_CTD_-PulM_CTD_ interface 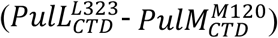, and 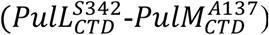. We used CSP as ambiguous restraints and co-evolution data as unambiguous restraints to generate the heterodimeric PulL_CTD_-PulM_CTD_ complex model. All 176 obtained models exhibit the same topology and the same assembly mode. In the best representative cluster with an ensemble of 53 models and a RMSD of 1.2 ± 0.9 Å, the interaction occurs between the respective homodimer interface 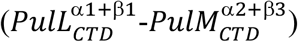 of each protein and the subunits are oriented in a parallel fashion (Figure 5G). The same orientation was seen in the heterodimer 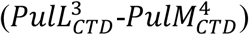 found in the crystal lattice and exhibiting the largest buried surface area, with a parallel orientation of protein subunits.

Interestingly, the same structural elements and the same residues are involved in the homodimer or heterodimer interfaces either through hydrophobic contacts (Figure 6A-D) or hydrogen bonds (Figure 6E-H). While PulM β3 and PulL β1 are each involved in an antiparallel β sheet formation within the homodimers (Figure 6E-G), they are hydrogen bonded together to form the heterodimer interface (Figure 6H). Additional hydrogen bonds involving 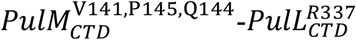 (in blue in Figure 6H) and 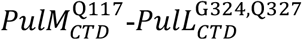 (in magenta in Figure 6H) further stabilize the heterodimer complex.

**Figure 6:**
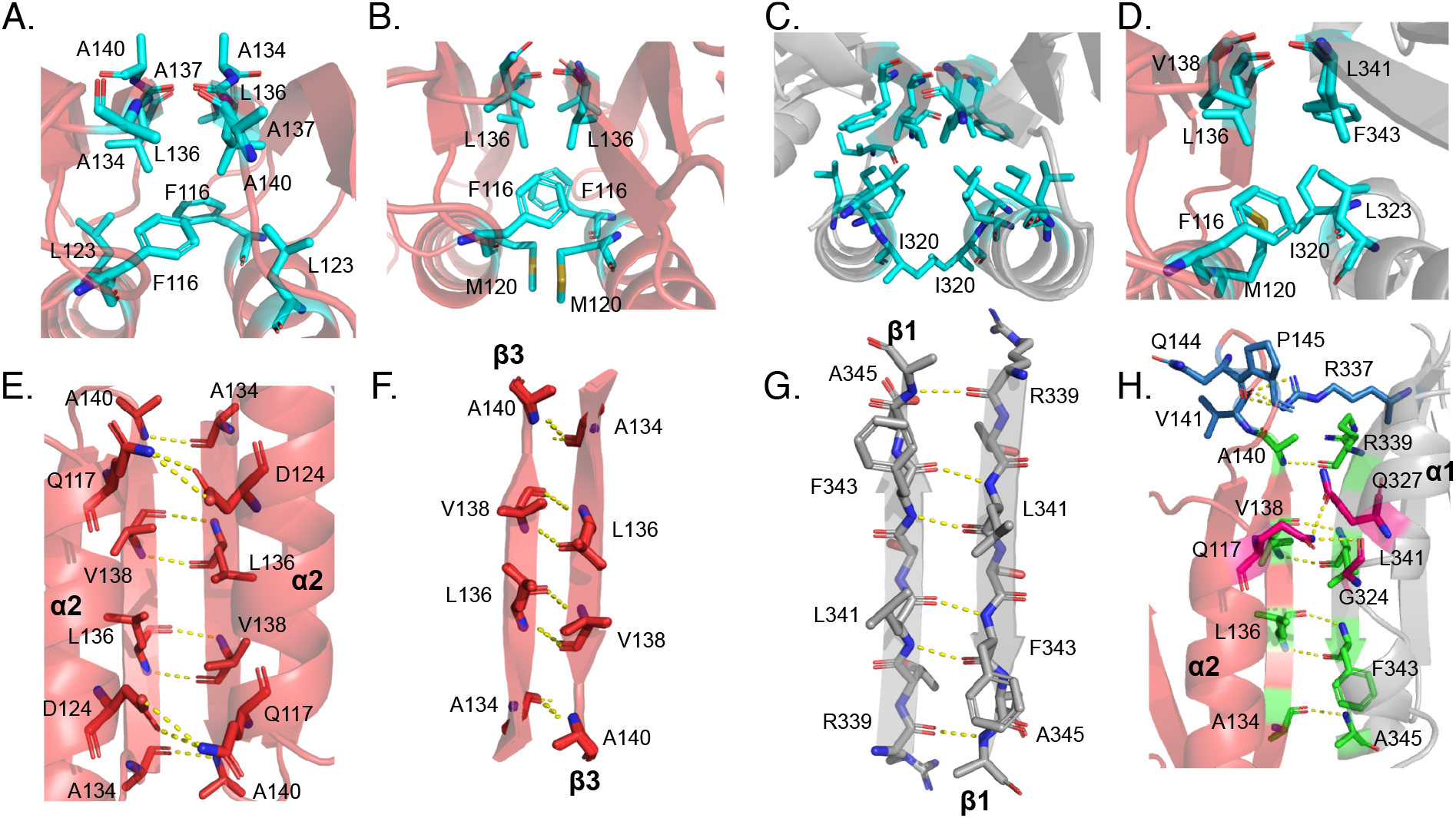
Comparison of the interfaces between PulL_CTD_ or PulM_CTD_ homodimers and the PulL_CTD_-PulM_CTD_ heterodimer complex. **A, B, C and D**. Hydrophobic interactions (cyan) in PulM_CTD_ (X-ray structure), PulM_CTD_ (NMR structure), PulL_CTD_, and the heterocomplex PulM_CTD_-PulL_CTD,_ from left to right. PulM_CTD_ is colored in red and PulL_CTD_ in grey. **E, F, G and H**. Hydrogen bonds (yellow dotted lines) in PulM_CTD_ (X-ray structure), PulM_CTD_ (NMR structure), PulL_CTD_, and the PulM_CTD_-PulL_CTD_ heterodimer complex, from left to right. In panel H, residues forming hydrogen bonds between β strands are shown in green, residues forming hydrogen bonds between α-helices in magenta and residues hydrogen bonding in the loops in blue. PulL and PulM protomers are colored in grey and red, respectively.

To evaluate the role of the PulL_CTD_-PulM_CTD_ association in protein secretion, we mutated the residues involved in the interface. We introduced single alanine substitutions in positions L319, L323, L341 and Q327 of PulL (Figure 7A, side chains shown as orange sticks) and tested the ability of these variants to restore pullulanase (PulA) secretion in *Escherichia coli* carrying the *pul* gene cluster with a non-polar *pulL* gene deletion in plasmid pCHAP8251. While Ala substitutions of Leu residues did not affect the function, replacing polar residue Q327 by an alanine led to a small but significant defect in PulA secretion (Figure 7B). This defect was further exacerbated in the presence of L319A or L341A substitutions (Figure 7B). On the PulM side, single Ala substitutions of F116, Q117, M120 and D124 of the α2 helix, or L136 of the β3 strand, did not affect the function (Figure 7C). However, the double substituted variant PulM^F116A,L136A^ showed a significant secretion defect, suggesting that the interface was weakened (Figure 7C). Replacing the surface exposed residue M120 by charged residues in PulM^M120D^ and PulM^M120K^ variants also significantly reduced secretion (Figure 7C). In comparison, charge inversions on the surface of the α1 helix, such as R88E and R92D, shown in cyan in Figure 7A, did not show any effect, consistent with their position distal to the interface with PulL.

**Figure 7.**
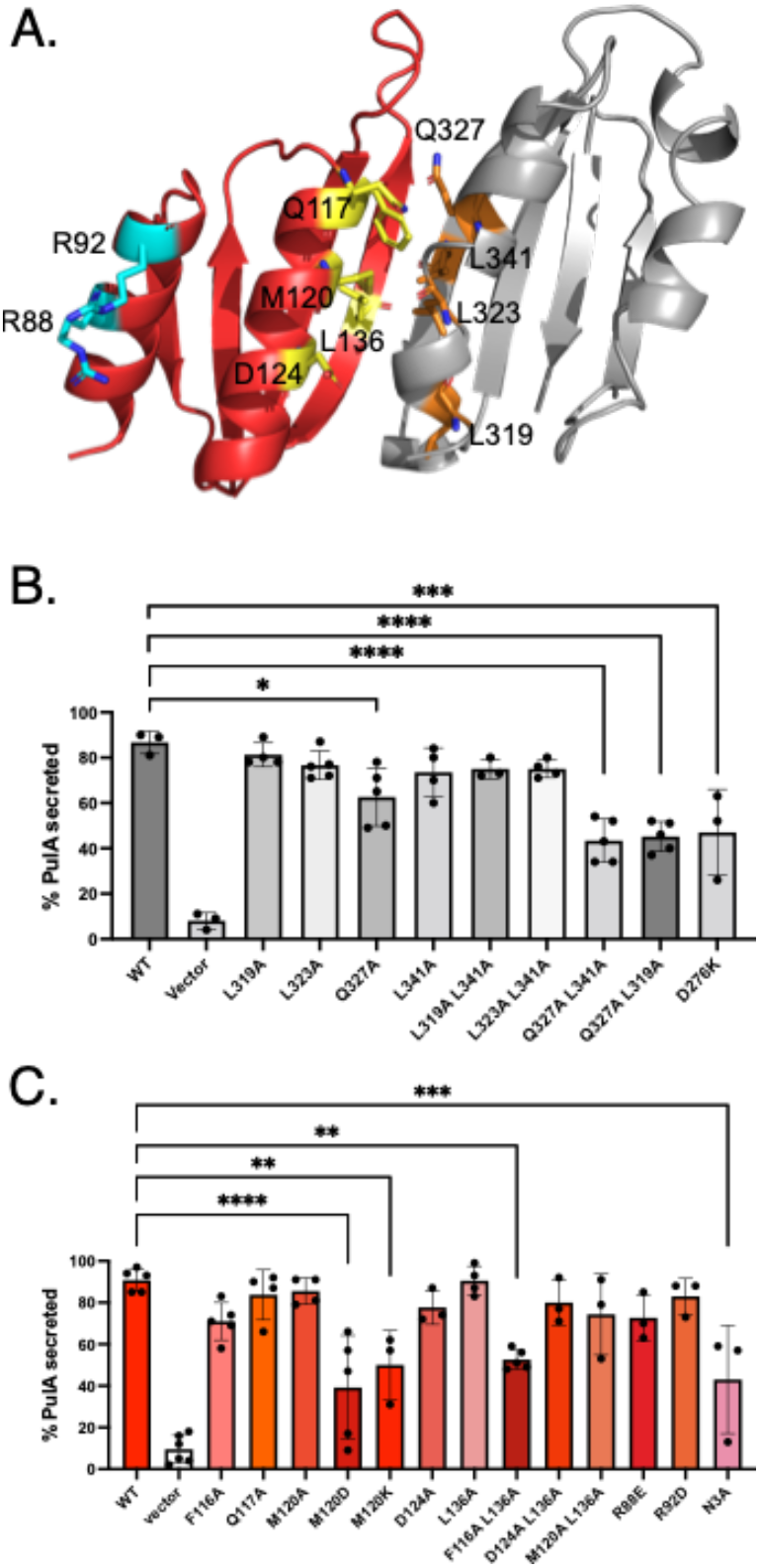
Functional characterization of the PulL_CTD_-PulM_CTD_ interface. **A**. Cartoon representation of the complex of PulL_CTD_ (grey) with PulM_CTD_ (red). The substituted residues are highlighted as sticks and marked with a single letter code, those at the PulL-PulM interfaces are colored in orange (PulL_CTD_) or yellow (PulM_CTD_), while those that are far from the interface are colored in cyan. **B**. PulA secretion in the presence of indicated PulL variants. Details of the secretion assays are provided in Materials and Methods. The bar graph heights represent the mean values and dots represent the percentage of secreted PulA from independent experiments. One-way ANOVA and multiple comparisons were done with GraphPad PRISM 9. Statistically significant differences relative to the wild type PulL are indicated. **C**. PulA secretion in the presence of indicated PulM variants. Data representation and analysis was performed as in (B).

Taken together, these data show that the intact PulL_CTD_-PulM_CTD_ interface and thus the PulL-PulM assembly plays an important role in the secretion. However, we cannot exclude that other PulL and PulM regions might also contribute to this interaction *in vivo*. One indication is the PulM^N3A^ variant mapping in the cytoplasmic N-terminus of the protein, which dramatically affected the stability of both proteins (Figure S7) and was defective in secretion (Figure 7C).

### Interactions of full-length membrane-anchored PulL and PulM

To gain further insights into the interaction between full-length PulL and PulM in the cellular context and in the presence of the membrane, we combined BACTH and cysteine cross-linking approaches. The BACTH approach allows us to analyze interactions between membrane proteins in their native environment (Karimova G *et al*., 1998). To study how the full-length PulL and PulM interact, we fused them to the C-terminal ends of T18 and T25, fragments of the *Bordetella pertussis* adenylyl cyclase (CyaA) catalytic domain. The resulting plasmids (listed in Table S3) were introduced in the *E. coli* strain DHT1 (Dautin *et al*., 2000) carrying a deletion of the endogenous *cya* gene. The CyaA T18 and T25 fragments do not interact and do not restore adenylyl cyclase activity as indicated by white colonies on indicator plates (Figure 8A) and low expression levels of the chromosomal *lacZ* gene (Figure 8B). Bacteria containing T18-PulL and T25-PulL hybrids also showed white colonies and low activity, comparable to the negative controls, indicating that the full-length membrane-embedded PulL does not form homodimers. Bacteria carrying the T18-PulM and T25-PulM chimera were pale blue on X-gal indicator and showed somewhat higher activity, indicating a tendency of PulM to homodimerize (Figure 8). Finally, bacteria co-producing PulL and PulM chimera showed strong and highly significant interaction signals. These results show that full-length PulL and PulM interact strongly with each other and preferentially form heterodimers *in vivo*.

**Figure 8.**
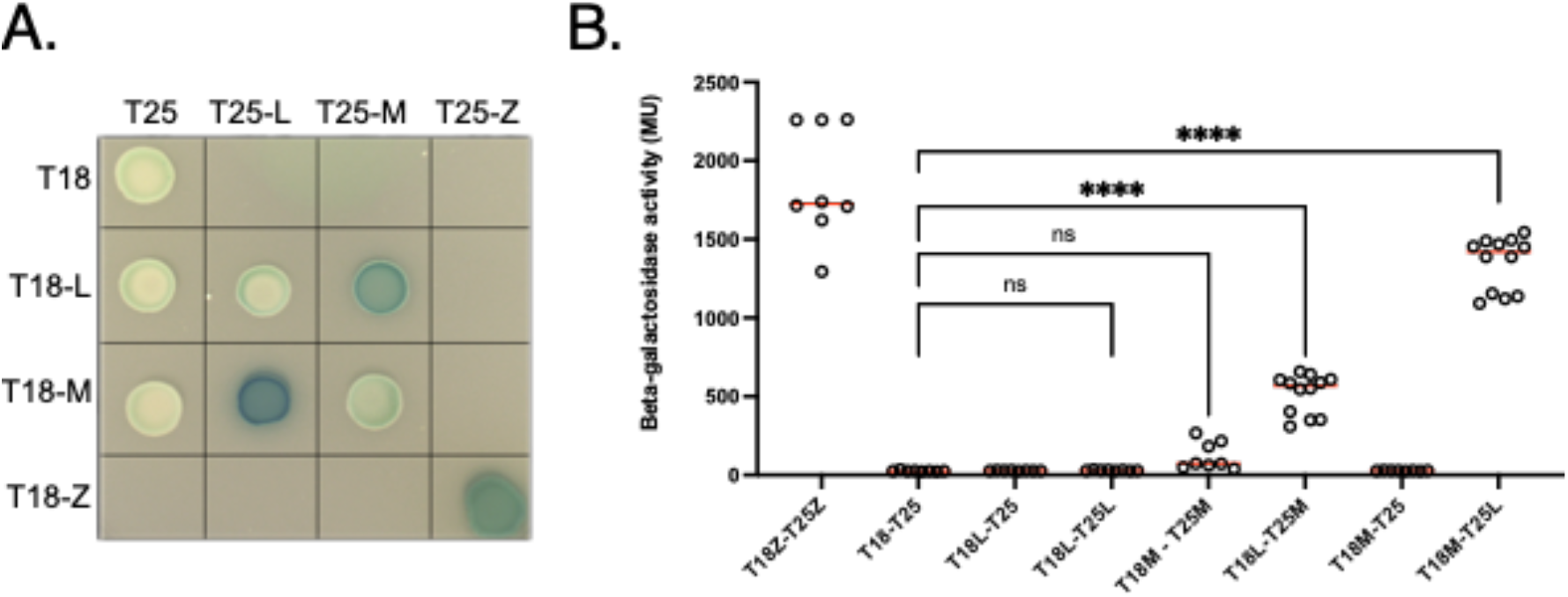
BACTH analysis of PulL and PulM interactions. A. Plate assay and B. Beta-galactosidase activity of strains producing T18, T25 or their chimera with yeast leucin Zipper (Z) (a positive control), PulL (L) or PulM (M). The β-galactosidase activities measured for independent bacterial cultures are plotted as dots and horizontal red lines show median values. Statistical analysis was performed with One-way ANOVA and multiple comparisons test, and plotted with GraphPad Prism 9 software.

PulL and PulM are type I membrane proteins with an N-in C-out orientation, each containing a single transmembrane (TM) segment. To investigate the involvement of these segments in the PulL-PulM interaction, we used cysteine crosslinking. Native PulM has one Cys residue in position 17, within its TM segment. We first replaced this residue by a Leu and used PulM^C17L^ as a starting construct to generate a series of PulM variants with single Cys substitution at positions 18 through 24. In a similar manner, we substituted Cys264, localized in the periplasmic part of PulL (PulL^C264L^) and used it to generate single Cys substitutions at positions 249 to 256 of the TM segment. Bacteria producing all the combinations of single Cys variants of PulL and PulM single Cys variants were treated with CuCl_2_ as an oxidant and their total extracts were analyzed by Western blot with anti-PulL or anti-PulM antibodies (Figure 9). The PulL^C264L^ variant did not produce any significant level of crosslinked species and a similar pattern was observed in the presence of single Cys residues at positions 249, 252, 253, 254, 255 and 256 (Figure 9). However, PulL^C264L^ with Cys at positions 250 and 251 produced some homodimers and also heterodimers, notably with PulM with Cys at positions 17 (PulM^WT^) and 18 (PulM^C17L-L18C^). The latter variant gave the most intense heterodimer bands, which were also detected with anti-PulM antibodies (Figure 9). The PulM variants with Cys at positions 17, 18 and 19 also produced homodimers bands. These results show that PulL and PulM can interact *via* their transmembrane segments and that in this region they also use the same interface for homo- and heterodimerization. The levels of PulM homodimers were not affected by the presence of PulL^I250C^ and PulL^V251C^, possibly because PulM was more abundant than PulL in these strains. Consistent with a higher fraction of PulL crosslinked with PulM, the L-M heterodimers were more readily detectable in anti-PulL Western blots compared to anti-PulM. Confirming the stabilizing role of PulM, the levels of all PulL variants were significantly reduced in the absence of PulM (Ø). In contrast, the levels of PulM did not change in the absence of PulL (Figure 9, last panel, Ø).

**Figure 9.**
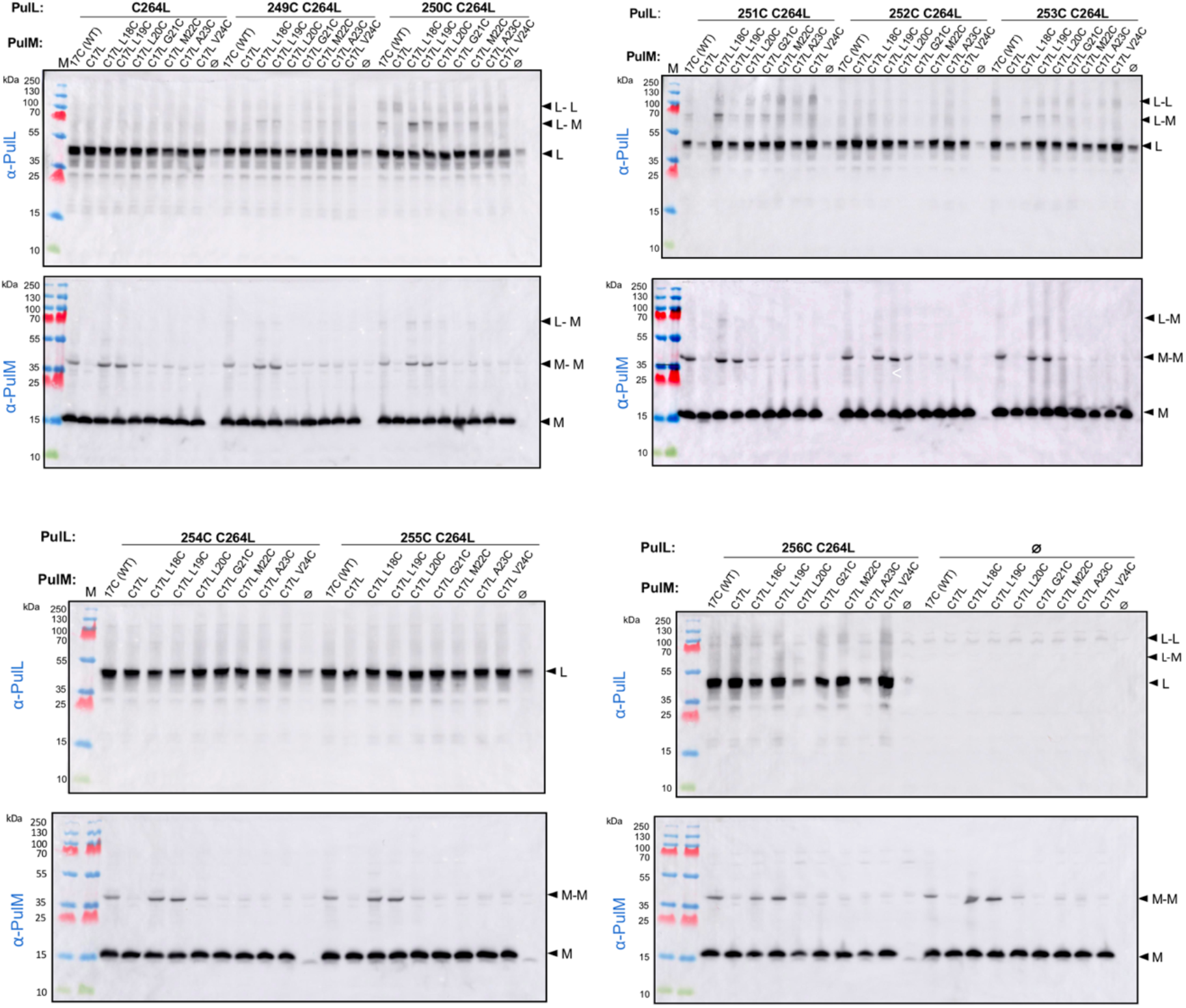
Cysteine scanning and crosslinking of PulL and PulM transmembrane segments. Bacteria producing the indicated PulL and PulM variants, or empty vector (Ø) were oxidized with CuCl_2_ and the total extract from 0.05 OD_600nm_ of bacteria was analyzed by SDS-PAGE and Western blot. Molecular weight markers are shown on the left, and migration of PulL (L) and PulM (M) monomers, homodimers (L-L, M-M) and heterodimers (L-M) are indicated on the right. The same total fractions were analyzed with anti-PulL antibody (top) and anti-PulM antibody (bottom).

The above experiments were performed in bacteria producing PulL and PulM in the absence of other T2SS components and at levels which may differ from those in the functional system. In the *pulC-O* operon encoding the *Klebsiella* T2SS, *pulL* and *pulM* genes are adjacent and transcribed from the same promoter. To determine their relative abundance, we quantified the PulL and PulM amounts in bacteria producing the functional T2SS from a single, moderate copy-number plasmid pCHAP8185 (Table S5). We used Western blot analysis with antibodies raised against PulL_CTD_ and PulM_CTD_ to quantify the level of these proteins in bacterial extracts expressing the T2SS genes and compared these signals with those known amounts of purified PulL_CTD_ and PulM_CTD_ (Figure S3). These measurements showed that PulM is present in large excess over PulL, with a molar ratio of about 20:1.

Combining the structural data on the PulL_CTD_-PulM_CTD_ heterodimer with the cysteine crosslinking experiments allowed us to build a 3D model of the membrane-anchored and periplasmic region of the PulL-PulM complex (Figure 10A and B). Sequence analysis revealed the presence of long periplasmic α-helices in both PulL and PulM, encompassing the transmembrane segments and upstream of the CTDs. Additionally, two regions involved in coiled-coil formation are predicted in both proteins, the first one overlapping with the transmembrane segments (Figure S8A-C). Consequently, periplasmic helices of PulL and PulM were modeled as a coiled-coil, in which knobs-into-holes packing was confirmed by SOCKET2 analysis (Kumar and Woolfson, 2021) (Figure S8D). In the model, the relative alignment of the predicted transmembrane segments is compatible with an orthogonal orientation of the connecting periplasmic helices with respect to the membrane. The coiled-coil formed by the periplasmic helices of PulL and PulM is followed by short disordered linkers connecting the α1 helices of the CTDs heterodimer. In the model of the complex, a potential salt-bridge is found between D276 of PulL and R42 of PulM (Figure 10B). Interestingly, the PulM^D276K^ variant displayed significantly reduced secretion (Figure 7B) which could indicate that the coiled-coil contributes to the overall interaction surface in the PulL-PulM complex.

**Figure 10:**
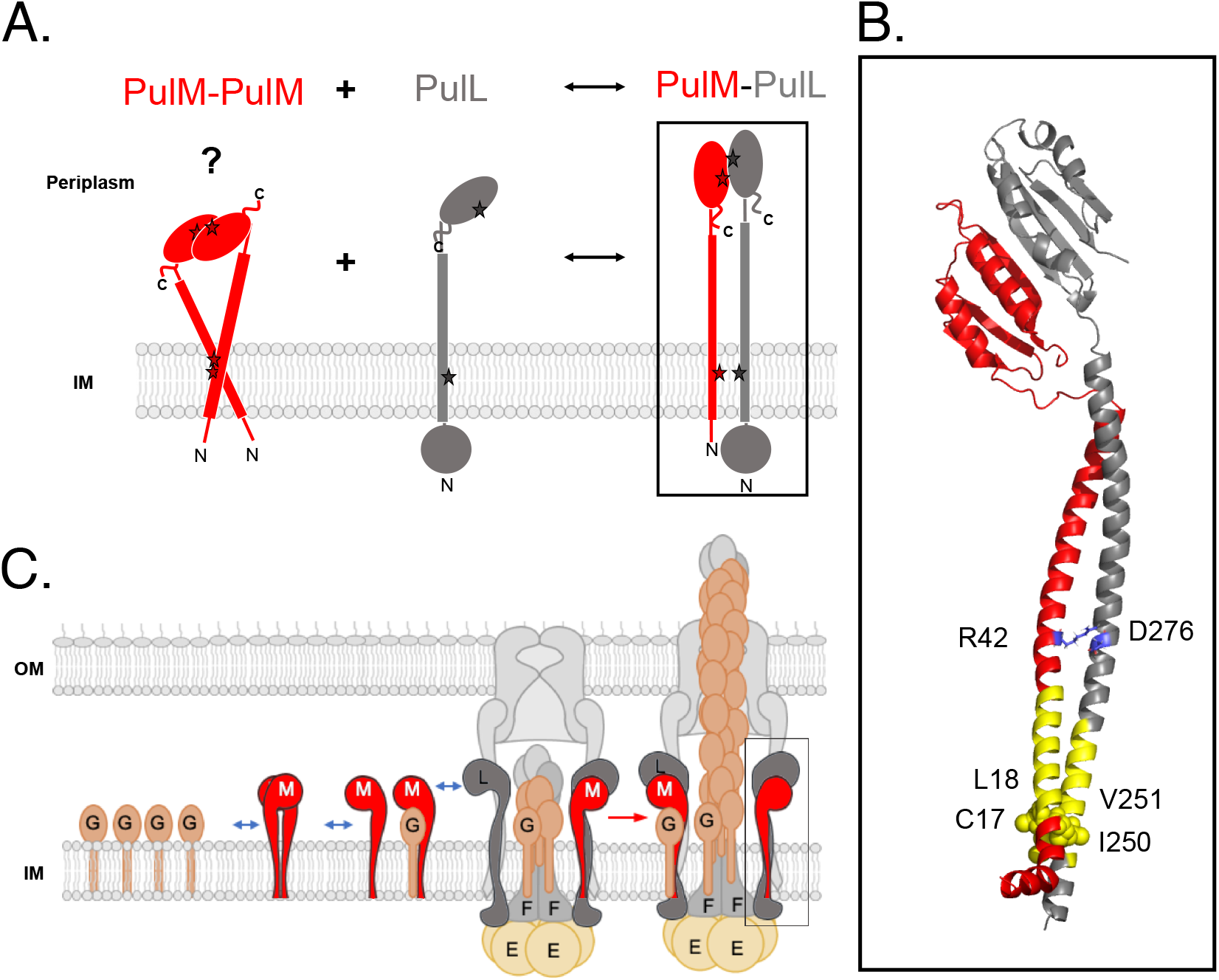
Full length PulL-PulM interaction model based on experimental results obtained in this study. **A**. Schematic view of heterodimerization process of PulL (grey) and PulM (red). PulM can behave as a homodimer (BACTH data) via interfaces at the C-terminal domains in the periplasm (NMR data) and in the inner membrane (IM) via their transmembrane domains (crosslinking data). PulL is in monomeric form (BACTH data). We postulate that PulL leads the PulL-PulM heterodimerization. The interface of heterodimerization is the same as the homodimerization interface (NMR and crosslinking data). Stars indicate region of interactions of PulL and PulM, in homo and heterodimeric forms. **B**. 3D model of the PulL-PulM complex based on the PulL_CTD_-PulM_CTD_ heterodimer structure (PDB ID: 8AB1) and from crosslinking data, both obtained in this study and on the structure of orthologue PilN-PilO complex (PDB ID: 3JC8). PulL is colored in grey, PulM in red, transmembrane segments in yellow. Residues close in space as revealed by crosslinking are annotated on the structural model. **C**. Model of full-length PulL and PulM in the context of the T2SS. Highly abundant PulG pseudopilus subunit interacts with PulM. By binding to PulL, PulM targets PulG to the PulL-PulE-PulF complex. PulG is released from PulM and is added to the growing pseudopilus.

## Discussion

Here we report the first structures of the *K. oxytoca* PulL_CTD_ and PulM_CTD_, and investigate their assembly in solution. By combining NMR and X-ray we show that while these two proteins form each a homodimer, they also assemble as a heterodimer. Intriguingly, the same interfaces and residues are involved in both homo- and heterodimerization, but resulting in different orientations of the subunits. These different parallel and antiparallel topologies also observed for other GspL, GspM and their orthologues indicate the high plasticity of their interfaces and their need for stabilization in a dimeric form (Figure S4, S5).

The ferredoxin-like fold of PulL_CTD_ and PulM_CTD_ and their orthologues explains well their structural complementarity and their dynamic assembly mode, both as homodimer and as heterodimer. The ferredoxin-like fold is highly favorable for molecular association. Its symmetrical repetition has been proposed by Eck and Dayhoff (known as Dayhoff’s theory) as the result of repetition of simple peptides to form a protein and would represent the protein ancestor (Eck and Dayhoff, 1966, Alva and Lupas, 2018). PulL_CTD_ and PulM_CTD_ indeed exhibit an internal pseudo-symmetry, where the two faces of the ferredoxin-like fold (α1-β1-β2 and α2-β3-β4) are structurally superimposable and present high level of sequence similarity (64% for PulM_CTD_ and 66 % for PulL_CTD_). Interestingly, the dimerization interfaces of PulL_CTD_ and PulM_CTD_ are also highly similar, at the sequence and structural levels, with overlapping residue positions involved in the interactions (Figure S6).

The PulL_CTD_ and PulM_CTD_ homodimers associate in an antiparallel fashion, whereas the PulL_CTD_-PulM_CTD_ association is parallel and thus more compatible with the membrane insertion of the proteins where both N-termini are found on the same side of the dimer (Figure 10). This interaction mode is consistent with the BACTH data wherein full-length membrane-inserted proteins form heterodimers in the parallel arrangement, and with previous data showing that the formation of GspL–GspM heterodimers in *D. dadantii* is favored over their homodimerization (Lallemand *et al*., 2013). Although the antiparallel homodimerization of PulL_CTD_ and PulM_CTD_ would require a tangled arrangement with the respect to the membrane (Figure 10A), we cannot exclude that it might occur during the process. The long coiled-coil regions and flexible linkers that connect them to the CTDs could provide additional plasticity in the periplasmic region allowing this antiparallel association.

In addition to the PulL_CTD_-PulM_CTD_ interface, native full-length forms of PulL and PulM also interact *via* their transmembrane segments, as shown by Cys scanning experiments. Like for the CTDs, we found an overlap between homo- and heterodimer interfaces involving the transmembrane segments (Figure 9). This is consistent with the previously observed competition between L-M homo- and heterodimerization (Lallemand *et al*., 2013). The relative abundance of these proteins *in vivo* is likely to determine whether homo- and heterodimers coexist and exchange. In most of the T2SS models, PulL and PulM are shown with a 1:1 stoichiometry, which finds confirmation in the purified T2SS subcomplex (Chernyatina and Low, 2019). However, in the cellular context and when the whole system is expressed, the amount of PulM is at least 20 times higher than that of PulL. Higher cellular levels of PulM compared to PulL have also been observed by fluorescence microscopy using the GFP (green fluorescent protein) chimera fused to their N-terminal ends (Buddelmeijer *et al*., 2006). Considering the static role of PulL, which serves as an anchor for the PulE ATPase, the higher abundance of PulM would ensure its dynamic role in PulG targeting (Nivaskumar *et al*., 2016, Santos-Moreno *et al*., 2017), which requires rapid exchange of its binding partners. The structural flexibility and complementarity of PulL_CTD_ and PulM_CTD_ is likely to facilitate this exchange during the highly dynamic pseudopilus assembly and turnover. The homodimerization would be more rapid than the heterodimerization that is more stable, similarly to Jun-Fos proteins where the homodimers are formed with a favorable kinetics, while the heterodimer is more stable, both forms are found *in vivo* and *in vitro* (Junius *et al*., 1996).

We have previously shown that PulG proteins that are assembled to form the pseudopilus (Lopez-Castilla *et al*., 2017), interact with PulM *via* their transmembrane parts (Nivaskumar *et al*., 2016, Santos-Moreno *et al*., 2017). By binding to PulL, PulM might therefore channel PulG to the assembly site defined by the PulE-PulL complex (Figure 10B). The higher abundance of PulM is consistent with its role in targeting of PulG to ensure its insertion into the growing pseudopilus. While PulG does not directly interact with PulL (Nivaskumar *et al*., 2016), crosslinking studies in *V. cholerae* showed their physical proximity (Gray *et al*., 2011) supporting the existence either of a ternary complex or their rapid exchange of partners. Rapid influx of pseudopilins would ensure efficient incorporation into helical filaments, thought to be the driving force for protein secretion. In addition to their role in pseudopilus assembly, the PulL and PulM homologues have been implicated in interactions with the secreted substrate in the *P. aeruginosa* T2SS (Michel-Souzy *et al*., 2018). Together, these data suggest that the PulL-PulM complex play a major role in the dynamic coupling of pseudopilus assembly and protein transport. Future studies are needed to address the influence of the pseudopilin and the secreted substrate on the AP assembly and to define the sequence of events during pseudopilin and substrate targeting to the secretion complex.

## Material and Methods

### Plasmid constructions

*Escherichia coli* K-12 strain DH5α F’*lacI*^*Q*^ was used as a host for cloning purposes. The plasmids used in this study are listed in Table S5. To construct plasmids pMS1222 and pMS1229, the *pulL* gene was PCR-amplified from plasmid pCHAP8258 as template using primers PulL Kpn 5 and PulL Eco 3 with the high-fidelity Q5 DNA polymerase (*New England Biolabs*). The PCR products were purified on a Qiaquick spin column, digested with *Eco*RI and *Kpn*I restriction enzymes (NEB) and ligated with the *Eco*RI - *Kpn*I digested plasmids pUT18C and pKT25. Site-directed mutagenesis to generate derivatives of pCHAP8258 and pCHAP1353 was performed using a two-step amplification with perfectly overlapping mutagenic primers. Two separate reactions with single mutagenic primers were performed for 6 cycles using the Q5 DNA polymerase under conditions recommended by the manufacturer. The reactions were mixed and additional 15 cycles of amplification were performed. The PCR reactions were treated with *Dpn*I and transformed into DH5α F’*lacI*^*Q*^ ultracompetent cells. The purified plasmids were verified by DNA sequencing *(Eurofins*). The list of oligonucleotides (*Eurofins*) is shown in Table S6.

Proteins PulL_CTD_ and PulM_CTD_ were produced in *E. coli* BL21 (DE3) under control of the T7 promoter. Bacteria were grown at indicated temperatures in LB medium or in minimal M9 medium (Miller, 1972). Antibiotics were added as required at following concentrations: ampicillin (Ap) 100 μg.mL^-1^, chloramphenicol (Cm), 25 μg.mL^-1^ and kanamycin (Km) at 25 μg.mL^-1^. When required, the genes under p*lacZ* control were induced with 1 mM isopropyl β-D-1-thiogalactopyranoside (IPTG).

### Production of unlabeled and isotope-labeled proteins

Derived pMalp2 vectors were used to express either the CTD of PulM (PulM_CTD_: from residues 79 to 161) or the CTD of PulL (PulL_CTD_: from residues 312 to 398) in *E. coli* BL21(DE3) cells (Table S5). Each protein was expressed in the periplasm fused to maltose binding protein (MBP) followed by a His_6_-tag and a TEV protease cleavage site that is used to remove the MBP and the His_6_-tag. Only an additional N-terminal serine remains after the cleavage.

Uniformly ^15^N and ^15^N/^13^C labeled PulM_CTD_, PulL_CTD_, were produced in M9 minimal medium using 1 g/L of ^15^NH_4_Cl and 4 g/L ^13^C glucose, as the sole nitrogen and carbon sources, respectively. Gene expression was induced with 1 mM IPTG (isopropyl β-D-1-thiogalactopyranoside) overnight at 18 °C in *E. coli* BL21 (DE3) cells. Unlabeled protein samples were prepared from *E. coli* BL21 (DE3) cell cultures in LB medium and expression was induced with 1 mM IPTG during 4 hours at 30°C.

### Protein purification

Proteins were purified from the supernatant after the sonication of bacterial cells and centrifugation at 16000 g, during 1 hour at 4°C. The supernatant of the cell lysis was filtered on a 0.22 μm filter, then loaded onto a HisTrap HP column (*Cytiva*) equilibrated with 50 mM Tris-HCl, pH 8.0, 100 mM NaCl, 10 mM imidazole. Bound proteins were eluted with a linear imidazole gradient going from 10 to 500 mM. The eluted fractions were then incubated with the TEV-His_6_ protease overnight at 10° C. The mixture was loaded on a HisTrap HP column to remove TEV-His_6_ and MBP-His_6_. Pul proteins from the unbound fractions were collected, concentrated on a 3 kDa cutoff centricon device (*Cytiva*) and applied on a Sephacryl S-100 column (*Cytiva)* equilibrated with 50 mM HEPES, pH 7.0, 50 mM NaCl. Fractions containing the purified Pul proteins were pooled and concentrated using centricon devices (*Cytiva*). Protease inhibitor cocktail, EDTA-free (*Roche*) was added to all buffers used during the purification. SDS-PAGE was used for analyzing the purity and protein contents of the fractions at each step of the purification.

PulM_CTD_ protein concentration was determined spectrophotometrically using the absorbance at 280 nm and a calculated extinction coefficient of 11000 M^-1^.cm^-1^. PulL_CTD_ concentration was obtained by BCA (BiCinchoninic acid Assay) method (Simpson, 2008), since this protein lacks tryptophane.

### NMR experiments for assignment

NMR spectra were acquired with a range of 0.3 to 0.4 mM ^15^N/^13^C labeled proteins in 50 mM HEPES, pH 7.0 (PulM_CTD_) or pH 6.5 (PulL_CTD_), 50 mM NaCl at 25°C on a 600 MHz Avance III HD and a 800 MHz Avance NEO spectrometers (*Bruker Biospin*) both equipped with a cryogenically cooled triple resonance ^1^H [^13^C /^15^N] probe (*Bruker Biospin*). The pulse sequences were employed as implemented in the TOPSPIN 3.6.1 (*Bruker, Biospin*) and IBS libraries (Favier and Brutscher, 2019). TOPSPIN 3.6.1 (*Bruker Biospin*) was used for NMR data acquisition and processing. The ^1^H, ^15^N, and ^13^C backbone and side chain resonance assignments were carried out as previously described (Dazzoni *et al*., 2021). Briefly standard experiments (Cavanagh *et al*., 1996) were used: 2D ^15^N-HSQC, ^13^C-HSQC, and 3D HNCA/HN(CO)CA, HNCACB/ HN(CO)CACB pair, HNCO/HN(CA)CO pair, HCCH-TOCSY, C(CO)NH-TOCSY and H(CCO)NH-TOCSY. For the assignment of C8H8 and CδHδ of phenylalanines and tyrosines, the 2D ^13^C-^1^H HBCBCGCDHD and HBCBCGCDCEHE spectra were used (Yamazaki *et al*., 1993). Side chain resonance assignments were completed by using 3D ^13^C and ^15^N NOESY-HSQC with mixing time of 120 ms optimized for either aliphatic or aromatic carbon detection (Iwahara *et al*., 2001), together with 2D ^1^H-^1^H NOESY (120 ms mixing time) and TOCSY in D_2_O.

2,2-Dimethyl-2-silapentane-5-sulfonate (DSS) signal was taken as 0 ppm for referencing proton chemical shifts and ^15^N and ^13^C chemical shifts were indirectly referenced to DSS (Wishart *et al*., 1995). CcpNmr Analysis (Vranken *et al*., 2005) was used for NMR data analysis. Analysis of the secondary structure was performed by using HN, Hα, Cα, Cβ, CO, and N chemical shifts with the TALOS-N prediction server (Shen and Bax, 2013).

PulM_CTD_ ^13^C/^15^N/^1^H resonances assignment were deposited to the BMRB under accession number 34719. PulL_CTD_ ^13^C/^15^N/^1^H resonances assignment were obtained in our previous work (Dazzoni *et al*., 2021).

### PulM_CTD_ and PulL_CTD_ NMR structure calculation

The structure of PulM_CTD_ and PulL_CTD_ in their dimeric and monomeric form respectively, were determined by performing several cycles of calculation with ARIA 2.3 software (Rieping *et al*., 2007) coupled to CNS 1.2 software and ARIAweb (Brünger *et al*., 1998, Allain *et al*., 2020), making use of the standard torsion angle simulating annealing protocol. Each cycle consisted of automatic 3D ^15^N-NOESY-HSQC and 3D ^13^C-NOESY-HSQC spectra assignment and structure calculations with 9 or 8 iterations with default parameters. In the last iteration 200 or 50 structures were calculated and further refined in an explicit water box (Linge *et al*., 2003). Some corrections to the NOE assignment were done manually. The 15 lowest energy structures exhibiting no NOE restraint violations > 0.5 Å and no dihedral angle violations > 5° were selected as the final ensemble.

For PulL_CTD_, an ensemble of 15 monomer structures was calculated. During the calculation process, most of the NOESs restraints from 3D ^15^N-NOESY-HSQC and 3D ^13^C-NOESY-HSQC spectra were automatically assigned by ARIA based on the chemical shifts previously obtained (Dazzoni *et al*., 2021).

PulM_CTD_ dimer structure was calculated in two steps, first by using intramolecular distance restraints derived from 3D ^13^C- and ^15^N-NOESY-HSQC spectra and the PulM_CTD_ ^13^C/^15^N/^1^H resonances assignment to determine the monomer structure. A 3D ^13^C/^15^N filtered NOESY-HSQC experiment (120 ms mixing time) was performed on a 1:1 double-labeled:unlabeled PulM_CTD_ mixture to obtain intermolecular distance restraints to calculate the dimer structure with ARIA. Chemical shift tolerances were set to 0.045 for protons and 0.4 ppm for the bound heteroatoms. Phi and psi dihedral angles were predicted with TALOS-N (Shen and Bax, 2013), and predictions classified as “strong” or “good” were incorporated as dihedral angle restraints. The structure ensemble was visualized and inspected with PyMOL (The PyMOL Molecular Graphics System, Version 2.0 Schrödinger, LLC), their quality was evaluated with PROCHECK-NMR (Laskowski *et al*., 1996) and the PSVS server (Bhattacharya *et al*., 2007). The atomic coordinates of PulM_CTD_ dimer and restraints used in the calculation were deposited in the Protein Data Bank (PDB ID: 7ZE0).

### Protein-protein interaction analysis by NMR

The protein-protein interactions were monitored by comparison of the ^1^H-^15^N HSQC spectra of one labeled protein alone and in the presence of its unlabeled partner at 25°C. ^1^H-^15^N HSQC experiments were acquired on 30 µM of either ^15^N-PulM_CTD_ or ^15^N-PulL_CTD_ alone or in the presence of 60 µM, 120 µM and 250 µM of unlabeled PulL_CTD_ or PulM_CTD_, respectively. To avoid dilution, the unlabeled protein was lyophilized in the same buffer and added to the labeled protein sample in solution. Chemical shift perturbations (CSP) of backbone amide cross-peaks were quantified by using the equation CSP = [ΔδH^2^ + (ΔδN*0.159)^2^]^1/2^, where ΔδH and ΔδN are the observed ^1^H and ^15^N chemical shift changes between the two experimental conditions. CSP higher than 1.5, 2 or 3 standard deviations (σ) from the mean were considered for the analysis.

### Native mass spectrometry

Prior to performing native mass analysis, the quality of each protein was assessed by intact mass measurement under denaturing conditions. For native mass spectrometry, protein samples were buffer exchanged against 250 mM ammonium acetate (pH 7.0) using Zeba spin desalting columns with a 7 kDa cutoff (*Thermo Fisher Scientific, Waltham, MA, USA*). The PulL_CTD_-PulM_CTD_ complex was either formed prior to or after buffer exchange by mixing PulM_CTD_ with an equimolar or a 2-fold molar excess PulL_CTD_.

Samples (at a final concentration of 5 to 10 µM) were analyzed on a SynaptG2-Si HDMS mass spectrometer (*Waters*) equipped with a nanoelectrospray source. The instrument was calibrated in sensitivity mode using a 2 mg/ml cesium iodide solution prepared in 50% isopropanol, 0.1% formic acid in the 50 to 5000 *m/z* range, and the quadrupole profile was adjusted to ensure the best transmission in the selected mass range. To preserve the integrity of non-covalent complexes in the gas phase, the instrument settings were carefully adjusted to the following values: capillary voltage, 1.5-2.0 kV; sampling cone, 150 V; source offset, 150 V; trap gas flow: between 5 and 7 mL/min, trap collision energy, 4 V; cone gas, 20 L/h; source temperature, 30°C. Spectra were acquired in positive mode for 5 to 10 minutes to obtain a good signal-to-noise ratio and processed with MassLynx 4.1 software (*Waters*) with minimal smoothing. The validation of the above-mentioned instrument settings was performed using myoglobin from equine skeletal muscle (*Sigma Aldrich*) prepared in 250 mM ammonium acetate buffer, pH 7.0 and using the same experimental procedure than for PulL_CTD_ and PulM_CTD_.

### Crystallization and diffraction data collection

For PulL_CTD_-PulM_CTD_ complex analysis, an equimolar mixture of two proteins was prepared at a final concentration of 1 mM and co-eluted on a Sephacryl S-100 column (*Cytiva)* equilibrated with 50 mM HEPES, pH 7.0, 50 mM NaCl. Initial screening of crystallization conditions was carried out by the vapor diffusion method using a MosquitoTM nanoliter-dispensing system (*TTP Labtech, Melbourn, United Kingdom*) following the established protocols (Weber *et al*., 2019). Briefly, sitting drops were set up using 400 nl of a 1:1 mixture of each sample protein and crystallization solutions (672 different commercially available conditions) equilibrated against a 150-μl reservoir in multiwell plates (*Greiner Bio-one, GmbH, Frichenhausen, Germany*). The crystallization plates were stored at 18°C in a RockImager1000^®^ (*Formulatrix, Bedford, MA, United States*) automated imaging system to monitor crystal growth. The best crystals were obtained in crystallization conditions containing 30%w/v PEG 4K, 0.1 M HEPES pH 7.5, 0.2 M CaCl_2_ for PulM_CTD_; 0.5 M LiSO_4_ and 15%w/v PEG 8K for PulL_CTD_; and 0%w/v PEG 3350, 0.2M KCl for the PulL_CTD_-PulM_CTD_ complex. Crystals were then flash cooled in liquid nitrogen using the condition of crystallization supplemented with 30% (V/V) of glycerol as cryoprotectant.

Diffraction data were collected at cryogenic temperatures (100k) on beamlines PROXIMA-1 and PROXIMA-2A at synchrotron SOLEIL (*St Aubin, France*) and processed with autoPROC (Vonrhein *et al*., 2011).

### X-ray structure determination and model refinement

The crystal structures of the PulL_CTD_, PulM_CTD_ and the PulL_CTD_-PulM_CTD_ complex were solved by the molecular replacement method with Phaser (McCoy *et al*., 2007), using NMR models as search probes. Final models were obtained through interactive cycles of manual model building with Coot (Emsley and Cowtan, 2004) and reciprocal space refinement with Buster (Bricogne *et al*., 2011) and REFMAC (Murshudov *et al*., 2011). X-ray diffraction data collection and model refinement statistics are summarized in Table S2. All structure figures were generated with Chimera (version 1.13rc) (Pettersen *et al*., 2004) or with PyMOL (version 2.5.2 The PyMOL Molecular Graphics System, Version 2.0 Schrödinger, LLC),)

### Protein secretion assays

Strain PAP7460 *Δ(lac-argF)U169 araD139 relA1 rpsL150 ΔmalE444 malG501 [F’ (lacI*^*Q*^ *ΔlacZM15 pro+ Tn10)] (Tc*^*R*^)) was used as a *pul* gene expression host (Possot *et al*., 2000). PAP7460 bacteria harboring plasmids pCHAP8251 or pCHAP8496 (Table S5) were transformed with compatible plasmids encoding PulL and PulM variants (Table S5). Bacteria were cultured overnight in LB containing Ap and Cm, 0.4% D-maltose and 0.1vol of M63 salts (Miller, 1972) at 30°C. The next day, 6 ml of the same fresh medium was inoculated with 300 μl of precultures and grown for 5 hours to OD_600nm_ > 1.8. Cultures were normalized of OD_600nm_ of 1 in a total volume of 1 mL and fractionated as follows: 0.1 mL of cultures were centrifuged for 10 min at 16000 x g in an Eppendorf centrifuge, the supernatant was aspirated off and the bacterial pellets were resuspended in 0.1 mL of SDS sample buffer to give the Cell fraction. The remaining 0.9 mL was centrifuged for 5 min at 16000 x g. The supernatant was transferred to a fresh tube and centrifuged for another 10 min at 16000 x g. A fraction (0.1 mL) of the supernatant was mixed with 0.1 mL of 2x SDS sample buffer to give the Supernatant fraction. The bacterial pellets of the 0.9 mL of normalized cultures were resuspended in 90 μL of SDS sample buffer for analysis of PulL and PulM levels (Figure S7). The Cell- and Supernatant fractions corresponding to the 0.05 OD_600nm_ of bacteria were analyzed on 10% Tris-glycine SDS-PAGE and transferred to the nitrocellulose membranes. The membranes were probed by Western blot with anti-PulA antibodies and revealed by fluorescence using ECL-2 and Typhoon SLA-9000. The concentrated cell pellets (0.1 OD_600nm_) were analyzed on 10% Tris-tricine SDS-PAGE and Western blot using anti-PulL and anti-PulM antibodies. Membranes were incubated with secondary anti-rabbit antibodies coupled to HRP (*Cytiva*) and revealed with ECL-2 (*Thermo*). The fluorescence signals were recorded using Typhoon SLA-9000. Signal intensity of PulA bands in cell- and supernatant fractions was quantified using ImageJ to calculate the fraction of PulA in the supernatant. Data were analyzed with Prism GraphPad software, using Ordinary one-way ANOVA test with multiple comparisons.

### Bacterial two-hybrid assays

Bacterial two-hybrid assays (Karimova G *et al*., 1998) were performed in strain DHT1 (Dautin *et al*., 2000). Plasmids encoding compatible pUT18C and pKT25 vectors and their derivatives were co-transformed in calcium-competent DHT1 bacteria and selected on LB plates containing Ap and Km. After 47-72 hours of growth at 30°C, single colonies were picked at random and inoculated in LB containing the same antibiotics. Bacteria were grown overnight at 30°C and the precultures were used to inoculate the same medium supplemented with 1 mM IPTG. After 4 hours, cultures were placed on ice and beta-galactosidase assays were performed as described previously (Miller, 1972). Beta-galactosidase activity from at least 8 independent cultures were plotted and the data was analysed using GraphPad Prism 9 software. For the qualitative plate tests, 10 μl of bacterial cultures was spotted on LB plates containing Ap, Km, X-gal (0.2 mg. mL^-1^) and 100 μM IPTG. The plates were cultured for 24-36 hours at 30°C and images were recorded with a digital camera.

### Cysteine crosslinking assays

Plasmids encoding different PulL and PulM variants or the empty vectors were co-transformed into strain PAP7460 and selected on LB Ap Cm plates at 30°C. Single colonies were inoculated into 5 mL of LB containing Ap and Cm and grown overnight at 30°C with shaking at 200 rpm. The next day, 0.2 mL of the precultures was diluted into 5 mL of fresh LB Ap Cm medium and bacteria were incubated for 5 hours at 30°C with vigorous shaking. Cultures were normalized to OD_600nm_ of 1 and 1 mL of bacteria was centrifuged for 3 min at 16000 x g in a table-top Eppendorf centrifuge. Bacterial pellets were washed once with phosphate buffer saline (PBS) and resuspended in 1 mL of buffer containing 50 mM MOPS pH 7.0, 5 mM MgCl_2_, 10% glycerol. Bacterial suspensions were prewarmed at 23°C, CuCl_2_ was added to 300 µM (final concentration) and incubated for 23 min at 23°C in a Thermomixer with shaking at 650 rpm. To stop the reaction, EDTA pH 8.0 was added to a final concentration of 22.5 mM. Reaction mixtures were pelleted by centrifugation and bacteria were resuspended in the SDS sample buffer at 10 OD_600nm_ mL^-1^. Total extracts were analyzed by sodium dodecyl sulfate gel electrophoresis (SDS-PAGE) on 10% Tris-Tricine gels. Proteins were transferred to nitrocellulose membranes (*Cytiva*) using the fast blotting system PowerBlot (Invitrogen) and a 1-Step transfer buffer (*Thermo*). The membranes were blocked in 5% skim milk in Tris buffer saline solution containing 0.05% Tween-20 (TBST) and probed with anti-PulL or anti-PulM antibodies. The signals were revealed with ECL chemiluminescence kit (*Thermo*) and recorded on Amersham 680 imager.

### Quantification of cellular PulL and PulM levels

To estimate the ratio between the cellular levels of PulL and PulM, we used semi-quantitative Western blot analysis. Different amounts of total bacterial extracts in parallel with the purified PulL_CTD_ or PulM_CTD_ were analyzed on 10% Tris-Tricine SDS-PAGE. Total bacterial extracts were prepared from cultures of strains PAP7460 harboring plasmid pCHAP8185 containing the *pul* operons. Bacteria were grown in LB supplemented with Ap, 0.4% D-maltose and 0.1 vol of M63 salts. Colony forming units from these cultures were counted by plating bacterial serial dilutions in triplicate. The proteins were transferred on nitrocellulose membranes, probed with antibodies directed against PulL_CTD_ and PulM_CTD_ domains followed by secondary goat anti-rabbit antibodies coupled with HRP. The blots were developed using the ECL-2 kit and quantified on Typhoon FLA-9000 imager. The bands were quantified with ImageJ. The standard curves of signal intensities as a function of known PulL_CTD_ or PulM_CTD_ molar concentrations were plotted and linear regression analysis was used to determine the amounts of PulL and PulM in total bacterial extracts, and their molar ratio, using the GraphPad Prism software.

### Full-length PulL-PulM heterocomplex modeling

To model the structure of the membrane embedded and periplasmic regions of the PulL/PulM complex, we used the structure of the PilN/PilO complex from the piliated state of the type IV pilus machine from *Myxococcus xanthus* (PDB ID: 3JC8) (Chang *et al*., 2016). First, the structure of PulL_CTD_-PulM_CTD_ heterodimer was superimposed onto the C-terminal domains (ferredoxin like-domains) of PilN and PilO. In both proteins, DeepCoil (Ludwiczak *et al*., 2019) predicts two regions potentially involved in coiled-coil formation. Thus, N-terminal helical region of PilN/PilO was used as template for comparative modeling of the homologous PulL/PulM region as a coiled-coil. The sequences of PulL and PulM encompassing the transmembrane segments (TMS) and the long helices between the TMS and the C-terminal domains, as predicted by PSIPRED (Jones, 1999), were aligned on the corresponding regions of PilN and PilO, respectively. In addition to sequence similarity, we consider the following to guide the alignment: i) the positions of the hydrophobic TMS, ii) the length of the predicted helices and iii) the expected spatial proximity of PulL residues 250/251 and PulM residues 17/18 from the cysteine cross-linking experiments. Using the resulting alignments (Figure S8C) and the PulL_CTD_-PulM_CTD_ heterodimer structure, a full atom model of PulL^239-398^-PulM^1-161^ was built by using Modeller (Sali and Blundell, 1993). SOCKET2 (Kumar and Woolfson, 2021) identified a coiled-coil interface involving helical segments from PulL (L254-R300) and PulM (G21-I67). Regions with knobs-into-holes packing between PulL and PulM correspond to the ones predicted as coiled-coils (Figure S8D).

### PulM_CTD_-PulL_CTD_ complex docking with HADDOCK

The High Ambiguity Driven DOCKing HADDOCK webserver v.2.4 was used to drive protein-protein dockings (van Zundert *et al*., 2016, Honorato *et al*., 2021). HADDOCK uses interaction data of partner molecules, which are introduced as ambiguous interaction restrictions to guide the docking process by active and passive residue selection. Active residues selected were based on CSP experimental data. For PulL_CTD_ residues chosen are the following: 319, 320, 321, 323, 326, 337 to 345. In the case of PulM_CTD_, active residues are: 117, 118, 119, 121, 124, 125, 127, 133, 134, 136. Passive residues were automatically selected around active residues by the server. To complete our data, we used as unambiguous restraints data given by EVcoupling (Hopf *et al*., 2014) that indicate two co-evolving pairs of residues located at the same interfaces (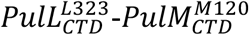, and 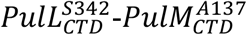). For those last restraints, the upper distance was set to 8.5 Å between the Cβ atoms of the involved residues. Input parameters for the HADDOCK server remained in the standard mode, with a minimum cluster size of 8. The 176 calculated models representing 88 % of the water-refined generated models, were clustered in 5 clusters. All clusters exhibit the same topologies, showing that based on the NMR and co-evolution data, only one assembly mode was possible. We considered cluster 2 as the best representative with a HADDOCK score of -79.9, and an ensemble of 53 structures with a RMSD of 1.2 ± 0.9 Å.

### Data deposition

Atomic coordinates and structure factors have been deposited in the RCSB Protein Data Bank under the accession codes 8A9W (PulL_CTD_), 7ZE0 (PulM_CTD_, NMR), 8A9X (PulM_CTD_, X-ray) and 8AB1 (PulL_CTD_-PulM_CTD_). PulM ^13^C/^15^N/^1^H resonance assignments were deposited to the BMRB under accession number 34719.

## Acknowledgements

This work was funded by the French Agence Nationale de la Recherche (ANR Synergy-T2SS ANR-19-CE11-0020-01) and the Fondation pour la Recherche Médicale (Equipe FRM 2017M.DEQ20170839114). YL was funded by the Pasteur Paris University (PPU) international PhD program and the China National Biotec Group Company Limited, and by a doctoral fellowship from the China Scholarship Council. We thank Ingrid Guilvout and Maylis Lejeune for their constant help. We acknowledge Iñaki Guijarro, Rémy Le Meur, Bertrand Raynal, Sébastien Brûlé and Christophe Thomas of C2RT for their help and assistance. The 800-MHz NMR spectrometer and the optima AUC of the Institut Pasteur were partially funded by the Région Ile de France (SESAME 2014 NMRCHR grant no 4014526) and DIM one health, respectively. The authors are grateful to the staff of the Institut Pasteur Crystallography platform for robot-driven crystallization screening. We acknowledge the Synchrotron SOLEIL (St Aubin, France) staff for assistance and advice during data collection on PROXIMA-1 and PROXIMA-2A beamlines. This work used the computational and storage service (TARS cluster) provided by the IT Department at Institut Pasteur, Paris.

## References

Abendroth, J., Kreger, A. & Hol, W. 2009. The dimer formed by the periplasmic domain of EpsL from the Type 2 Secretion System of Vibrio parahaemolyticus. Journal of structural biology, 168.

Abendroth, J., Murphy, P., Sandkvist, M., Bagdasarian, M. & Hol, W. 2005. The X-ray structure of the type II secretion system complex formed by the N-terminal domain of EpsE and the cytoplasmic domain of EpsL of Vibrio cholerae. Journal of molecular biology, 348.

Allain, F., Mareuil, F., Ménager, H., Nilges, M. & Bardiaux, B. 2020. ARIAweb: a server for automated NMR structure calculation. Nucleic acids research, 48.

Alva, V. & Lupas, A. 2018. From ancestral peptides to designed proteins. Current opinion in structural biology, 48.

Berry, J. & Pelicic, V. 2015. Exceptionally widespread nanomachines composed of type IV pilins: the prokaryotic Swiss Army knives. FEMS microbiology reviews, 39.

Bhattacharya, A., Tejero, R. & Montelione, G. 2007. Evaluating protein structures determined by structural genomics consortia. Proteins, 66.

Bricogne, G., Blanc, E., Brandl, M., Flensburg, C., Keller, P., Paciorek, W., Roversi, P., Sharff, A., Smart, O. S., Vonrhein, C. & Womack, T. O. 2011. BUSTER version 2.11.1. Cambridge, UK: Global Phasing Ltd.

Brünger, A. T., Adams, P. D., Clore, G. M., Delano, W. L., Gros, P., Grosse-Kunstleve, R. W., Jiang, J. S., Kuszewski, J., Nilges, M., Pannu, N. S., Read, R. J., Rice, L. M., Simonson, T. & Warren, G. L. 1998. Crystallography & NMR system: A new software suite for macromolecular structure determination. Acta crystallographica. Section D, Biological crystallography, 54.

Buddelmeijer, N., Francetic, O. & Pugsley, A. 2006. Green fluorescent chimeras indicate nonpolar localization of pullulanase secreton components PulL and PulM. Journal of bacteriology, 188.

Cavanagh, J., Fairbrother, W. J., Palmer III, A. G. & Skelton, N. J. 1996. Protein NMR Spectroscopy: Principles and Practice, New York, NY, U.S.A.

Chang, Y., Rettberg, L., Treuner-Lange, A., Iwasa, J., Søgaard-Andersen, L. & Jensen, G. 2016. Architecture of the type IVa pilus machine. Science (New York, N.Y.), 351.

Chernyatina, A. & Low, H. 2019. Core architecture of a bacterial type II secretion system. Nature communications, 10.

Cianciotto, N. P. & White, R. C. 2017. Expanding Role of Type II Secretion in Bacterial Pathogenesis and Beyond. Infection and immunity, 85.

D’enfert, C., Ryter, A. & Pugsley, A. P. 1987. Cloning and expression in Escherichia coli of the Klebsiella pneumoniae genes for production, surface localization and secretion of the lipoprotein pullulanase. The EMBO journal, 6.

Dautin, N., Karimova, G., Ullmann, A. & Ladant, D. 2000. Sensitive genetic screen for protease activity based on a cyclic AMP signaling cascade in Escherichia coli. Journal of bacteriology, 182.

Dazzoni, R., LóPez-Castilla, A., Cordier, F., Bardiaux, B., Nilges, M., Francetic, O. & Izadi-Pruneyre, N. 2021. 1 H, 15 N and 13 C resonance assignments of the C-terminal domain of PulL, a component of the Klebsiella oxytoca type II secretion system. Biomolecular NMR assignments.

Denise, R., Abby, S. & Rocha, E. 2019. Diversification of the type IV filament superfamily into machines for adhesion, protein secretion, DNA uptake, and motility. PLoS biology, 17.

Eck, R. & Dayhoff, M. 1966. Evolution of the structure of ferredoxin based on living relics of primitive amino Acid sequences. Science (New York, N.Y.), 152.

Emsley, P. & Cowtan, K. 2004. Coot: model-building tools for molecular graphics. Acta crystallographica. Section D, Biological crystallography, 60.

Favier, A. & Brutscher, B. 2019. NMRlib: user-friendly pulse sequence tools for Bruker NMR spectrometers. Journal of biomolecular NMR, 73.

Fulara, A., Vandenberghe, I., Read, R. J., Devreese, B. & Savvides, S. N. 2018. Structure and oligomerization of the periplasmic domain of GspL from the type II secretion system of Pseudomonas aeruginosa. Sci Rep, 8, 16760.

Ghosal, D., Kim, K. W., Zheng, H., Kaplan, M., Truchan, H. K., Lopez, A. E., Mcintire, I. E., Vogel, J. P., Cianciotto, N. P. & Jensen, G. J. 2019. In vivo structure of the Legionella type II secretion system by electron cryotomography. Nat Microbiol, 4, 2101–2108.

Gray, M. D., Bagdasarian, M., Hol, W. G. J. & Sandkvist, M. 2011. In vivo cross-linking of EpsG to EpsL suggests a role for EpsL as an ATPase-pseudopilin coupling protein in the Type II secretion system of Vibrio cholerae. Mol Microbiol., 79, 786–798.

Hobbs, M. & Mattick, J. 1993. Common components in the assembly of type 4 fimbriae, DNA transfer systems, filamentous phage and protein-secretion apparatus: a general system for the formation of surface-associated protein complexes. Molecular microbiology, 10.

Honorato, R., Koukos, P., Jiménez-García, B., Tsaregorodtsev, A., Verlato, M., Giachetti, A., Rosato, A. & Bonvin, A. 2021. Structural Biology in the Clouds: The WeNMR-EOSC Ecosystem. Frontiers in molecular biosciences, 8.

Hopf, T., Schärfe, C., Rodrigues, J., Green, A., Kohlbacher, O., Sander, C., Bonvin, A. & Marks, D. 2014. Sequence co-evolution gives 3D contacts and structures of protein complexes. eLife, 3.

Iwahara, J., Wojciak, J. M. & Clubb, R. T. 2001. Improved NMR spectra of a protein-DNA complex through rational mutagenesis and the application of a sensitivity optimized isotope-filtered NOESY experiment. Journal of biomolecular NMR, 19.

Janin, J., Rodier, F., Chakrabarti, P. & Bahadur, R. 2007. Macromolecular recognition in the Protein Data Bank. Acta crystallographica. Section D, Biological crystallography, 63.

Jiménez-García, B., Elez, K., Koukos, P., Bonvin, A. & Vangone, A. 2019. PRODIGY-crystal: a web-tool for classification of biological interfaces in protein complexes. Bioinformatics (Oxford, England), 35.

Jones, D. 1999. Protein secondary structure prediction based on position-specific scoring matrices. Journal of molecular biology, 292.

Junius, F., O’donoghue, S., Nilges, M., Weiss, A. & King, G. 1996. High resolution NMR solution structure of the leucine zipper domain of the c-Jun homodimer. The Journal of biological chemistry, 271.

Karimova G, Pidoux J, Ullmann A & Ladant D 1998. A bacterial two-hybrid system based on a reconstituted signal transduction pathway. Proceedings of the National Academy of Sciences of the United States of America, 95.

Krissinel, E. & Henrick, K. 2007. Inference of macromolecular assemblies from crystalline state. J. Mol. Biol., 372, 774–797.

Kumar, P. & Woolfson, D. 2021. Socket2: A Program for Locating, Visualising, and Analysing Coiled-coil Interfaces in Protein Structures. Bioinformatics (Oxford, England), 37.

Lallemand, M., Login, F. H., Guschinskaya, N., Pineau, C., Effantin, G., Robert, X. & Shevchik, V. E. 2013. Dynamic interplay between the periplasmic and transmembrane domains of GspL and GspM in the type II secretion system. PLoS One, 8, e79562.

Laskowski, R., Rullmannn, J., Macarthur, M., Kaptein, R. & Thornton, J. 1996. AQUA and PROCHECK-NMR: programs for checking the quality of protein structures solved by NMR. Journal of biomolecular NMR, 8.

Leighton, T., Mok, M., Junop, M., Howell, P. & Burrows, L. 2018. Conserved, unstructured regions in Pseudomonas aeruginosa PilO are important for type IVa pilus function. Scientific reports, 8.

Leighton, T., Yong, D., Howell, P. & Burrows, L. 2016. Type IV Pilus Alignment Subcomplex Proteins PilN and PilO Form Homo- and Heterodimers in Vivo. The Journal of biological chemistry, 291.

Linge, J., Williams, M., Spronk, C., Bonvin, A. & Nilges, M. 2003. Refinement of protein structures in explicit solvent. Proteins, 50.

Lopez-Castilla, A., Thomassin, J. L., Bardiaux, B., Zheng, W., Nivaskumar, M., Yu, X., Nilges, M., Egelman, E. H., Izadi-Pruneyre, N. & Francetic, O. 2017. Structure of the calcium-dependent type 2 secretion pseudopilus. Nat Microbiol, 2, 1686–1695.

Ludwiczak, J., Winski, A., Szczepaniak, K., Alva, V. & Dunin-Horkawicz, S. 2019. DeepCoil-a fast and accurate prediction of coiled-coil domains in protein sequences. Bioinformatics (Oxford, England), 35.

Luna Rico, A., Zheng, W., Petiot, N., Egelman, E. & Francetic, O. 2019. Functional reconstitution of the type IVa pilus assembly system from enterohaemorrhagic Escherichia coli. Molecular microbiology, 111.

Maffei, B., Francetic, O. & Subtil, A. 2017. Tracking Proteins Secreted by Bacteria: What’s in the Toolbox? Frontiers in cellular and infection microbiology, 7.

Mccoy, A., Grosse-Kunstleve, R., Adams, P., Winn, M., Storoni, L. & Read, R. 2007. Phaser crystallographic software. Journal of applied crystallography, 40.

Michel-Souzy, S., Douzi, B., Cadoret, F., Raynaud, C., Quinton, L., Ball, G. & Voulhoux, R. 2018. Direct interactions between the secreted effector and the T2SS components GspL and GspM reveal a new effector-sensing step during type 2 secretion. J Biol Chem, 293, 19441–19450.

Miller, J. H. 1972. Experiments in Molecular Genetics, New York, Cold Spring Harbor.

Murshudov, G., Skubák, P., Lebedev, A., Pannu, N., Steiner, R., Nicholls, R., Winn, M., Long, F. & Vagin, A. 2011. REFMAC5 for the refinement of macromolecular crystal structures. Acta crystallographica. Section D, Biological crystallography, 67.

Naskar, S., Hohl, M., Tassinari, M. & Low, H. 2021. The structure and mechanism of the bacterial type II secretion system. Molecular microbiology, 115.

Nivaskumar, M., Santos-Moreno, J., Malosse, C., Nadeau, N., Chamot-Rooke, J., Tran Van Nhieu, G. & Francetic, O. 2016. Pseudopilin residue E5 is essential for recruitment by the type 2 secretion system assembly platform. Mol Microbiol, 101, 924–41.

Peabody, C., Chung, Y., Yen, M., Vidal-Ingigliardi, D., Pugsley, A. & Saier, M. 2003. Type II protein secretion and its relationship to bacterial type IV pili and archaeal flagella. Microbiology (Reading, England), 149.

Pettersen, E., Goddard, T., Huang, C., Couch, G., Greenblatt, D., Meng, E. & Ferrin, T. 2004. UCSF Chimera--a visualization system for exploratory research and analysis. Journal of computational chemistry, 25.

Possot, O. M., Vignon, G., Bomchil, N., Ebel, F. & Pugsley, A. P. 2000. Multiple interactions between pullulanase secreton components involved in stabilization and cytoplasmic membrane association of PulE. J Bacteriol, 182, 2142–52.

Pugsley, A. P. 1993. The complete general secretory pathway in gram-negative bacteria. Microbiological reviews, 57.

Py, B., Loiseau, L. & Barras, F. 2001. An inner membrane platform in the type II secretion machinery of Gram-negative bacteria. EMBO reports, 2.

Rieping, W., Habeck, M., Bardiaux, B., Bernard, A., Malliavin, T. & Nilges, M. 2007. ARIA2: automated NOE assignment and data integration in NMR structure calculation. Bioinformatics (Oxford, England), 23.

Sali, A. & Blundell, T. 1993. Comparative protein modelling by satisfaction of spatial restraints. Journal of molecular biology, 234.

Sampaleanu, L., Bonanno, J., Ayers, M., Koo, J., Tammam, S., Burley, S., Almo, S., Burrows, L. & Howell, P. 2009. Periplasmic domains of Pseudomonas aeruginosa PilN and PilO form a stable heterodimeric complex. Journal of molecular biology, 394.

Sandkvist, M., Hough, L. P., Bagdasarian, M. M. & Bagdasarian, M. 1999. Direct interaction of the EpsL and EpsM proteins of the general secretion apparatus in Vibrio cholerae. J Bacteriol, 181, 3129–35.

Santos-Moreno, J., East, A., Guilvout, I., Nadeau, N., Bond, P. J., Tran Van Nhieu, G. & Francetic, O. 2017. Polar N-terminal Residues Conserved in Type 2 Secretion Pseudopilins Determine Subunit Targeting and Membrane Extraction Steps during Fibre Assembly. Journal of molecular biology, 429.

Shen, Y. & Bax, A. 2013. Protein backbone and sidechain torsion angles predicted from NMR chemical shifts using artificial neural networks. Journal of biomolecular NMR, 56.

Simpson, R. 2008. Quantifying protein by bicinchoninic Acid. CSH protocols, 2008.

Van Zundert, G., Rodrigues, J., Trellet, M., Schmitz, C., Kastritis, P., Karaca, E., Melquiond, A., Van, D. M, De Vries, S. & Bonvin, A. 2016. The HADDOCK2.2 Web Server: User-Friendly Integrative Modeling of Biomolecular Complexes. Journal of molecular biology, 428.

Vonrhein, C., Flensburg, C., Keller, P., Sharff, A., Smart, O., Paciorek, W., Womack, T. & Bricogne, G. 2011. Data processing and analysis with the autoPROC toolbox. Acta crystallographica. Section D, Biological crystallography, 67.

Vranken, W. F., Boucher, W., Stevens, T. J., Fogh, R. H., Pajon, A., Llinas, M., Ulrich, E. L., Markley, J. L., Ionides, J. & Laue, E. D. 2005. The CCPN data model for NMR spectroscopy: development of a software pipeline. Proteins, 59.

Weber, P., Pissis, C., Navaza, R., Mechaly, A., Saul, F., Alzari, P. & Haouz, A. 2019. High-Throughput Crystallization Pipeline at the Crystallography Core Facility of the Institut Pasteur. Molecules (Basel, Switzerland), 24.

Wishart, D., Bigam, C., Yao, J., Abildgaard, F., Dyson, H., Oldfield, E., Markley, J. & Sykes, B. 1995. 1H, 13C and 15N chemical shift referencing in biomolecular NMR. Journal of biomolecular NMR, 6.

Yamazaki, T., Yoshida, M. & Nagayama, K. 1993. Complete assignments of magnetic resonances of ribonuclease H from Escherichia coli by double- and triple-resonance 2D and 3D NMR spectroscopies. Biochemistry, 32.

Yueh, C., Hall, D., Xia, B., Padhorny, D., Kozakov, D. & Vajda, S. 2017. ClusPro-DC: Dimer Classification by the Cluspro Server for Protein-Protein Docking. Journal of molecular biology, 429.

