## Supplementary material for "Structure and dynamic association of an assembly platform subcomplex of the bacterial type II secretion system"

**Supplementary data: Oligomeric state of PulL$_{CTD}$ and PulM$_{CTD}$**
**Supplementary result**
**Supplementary Figure S1: Oligomeric state of PulL$_{CTD}$, PulM$_{CTD}$ and PulL$_{CTD}$-PulM$_{CTD}$ complex analysis.**
**Supplementary Figure S2:** Superposition of PulM$_{CTD}$ X-ray structure with PulM$_{CTD}$ NMR structure.
**Supplementary Figure S3:** Estimating the PulL and PulM molar ratio in the bacteria.
**Supplementary Figure S4:** Homologues and orthologues of PulL
**Supplementary Figure S5:** Homologues and orthologues of PulM
**Supplementary Figure S6:** Structural alignments of PulM$_{CTD}$ and PulL$_{CTD}$ dimerization regions.
**Supplementary Figure S7:** Levels of PulL and PulM variants in bacterial extracts.
**Supplementary Figure S8:** Sequence and structure analysis used for PulL-PulM complex modeling.

**Supplementary Table S1:** PulL$_{CTD}$ monomer NMR structure statistics and restraints
**Supplementary Table S2:** Crystallography data collection and refinement statistics

**Supplementary Table S3:** $PulM_{CTD}$ dimer NMR structure statistics and restraints

**Supplementary Table S4:** Scoring biological interfaces in the X-ray crystallographic structure of the $PulL_{CTD}$-$PulM_{CTD}$ heterodimer complex

**Supplementary Table S5**: Plasmids used in this study

**Supplementary Table S6:** Oligonucleotides used in this study

**Supplementary Materials and Methods**

**Supplementary References**

**Supplementary data: Oligomeric state of PulL$_{CTD}$ and PulM$_{CTD}$**

Oligomeric forms of both PulL$_{CTD}$ and PulM$_{CTD}$ have been first suspected by NMR analysis. The rotational correlation time ($\tau_c$) was estimated as 10.8 ns for PulL$_{CTD}$ and 8.90 ns for PulM$_{CTD}$ by using $T_1$ and $T_2$ relaxation times measurements NMR experiments. For both proteins, these values are consistent with a dimeric form with a molecular weight of 19.07 kDa for PulL$_{CTD}$ and 17.86 kDa PulM$_{CTD}$, rather than a monomeric form (Figure S1A). To determine the oligomeric states more accurately, we performed analytical ultracentrifugation (AUC). As the NMR experimental conditions require high protein concentrations (300-400 μM in our experiments), we performed AUC with samples at concentrations ranging from 10 to 300 μM.

For PulL$_{CTD}$, we observed two main peaks with sedimentation coefficient of 1 and 1.81 S (Figure S1C). These two peaks correspond to a monomeric and a dimeric form. We observed an additional peak with higher sedimentation coefficient, compatible with a tetrameric form that evolves to a larger oligomeric form at high concentration. Nevertheless, the dimeric form represents the major form in solution (≅75% of total protein) even at low concentrations, compared to the monomeric species. A monomer-dimer equilibrium can thus exist depending on the protein concentration. Evidence for this exchange at the NMR time scale (μs-ms) is observed on the PulL$_{CTD}$ $^{1}H$-$^{15}N$ HSQC spectrum (Figure 1A) where the line widths were broader than would be expected for a monomeric protein.

Regarding PulM$_{CTD}$, we obtained a sedimentation coefficient of 1.6 S, (Figure S1D) for all tested concentrations, including that of the NMR sample (300 μM). These values are compatible with a dimeric form of PulM$_{CTD}$. Nonetheless, the PulM$_{CTD}$ peaks are quite broad at the tested concentrations (Figure S1D) and are getting narrower when increasing concentrations, suggesting a monomer-dimer equilibrium.

**Supplementary Figure S1: Oligomeric state of $PulL_{CTD}$, $PulM_{CTD}$ and $PulL_{CTD}$-$PulM_{CTD}$ complex analysis.**

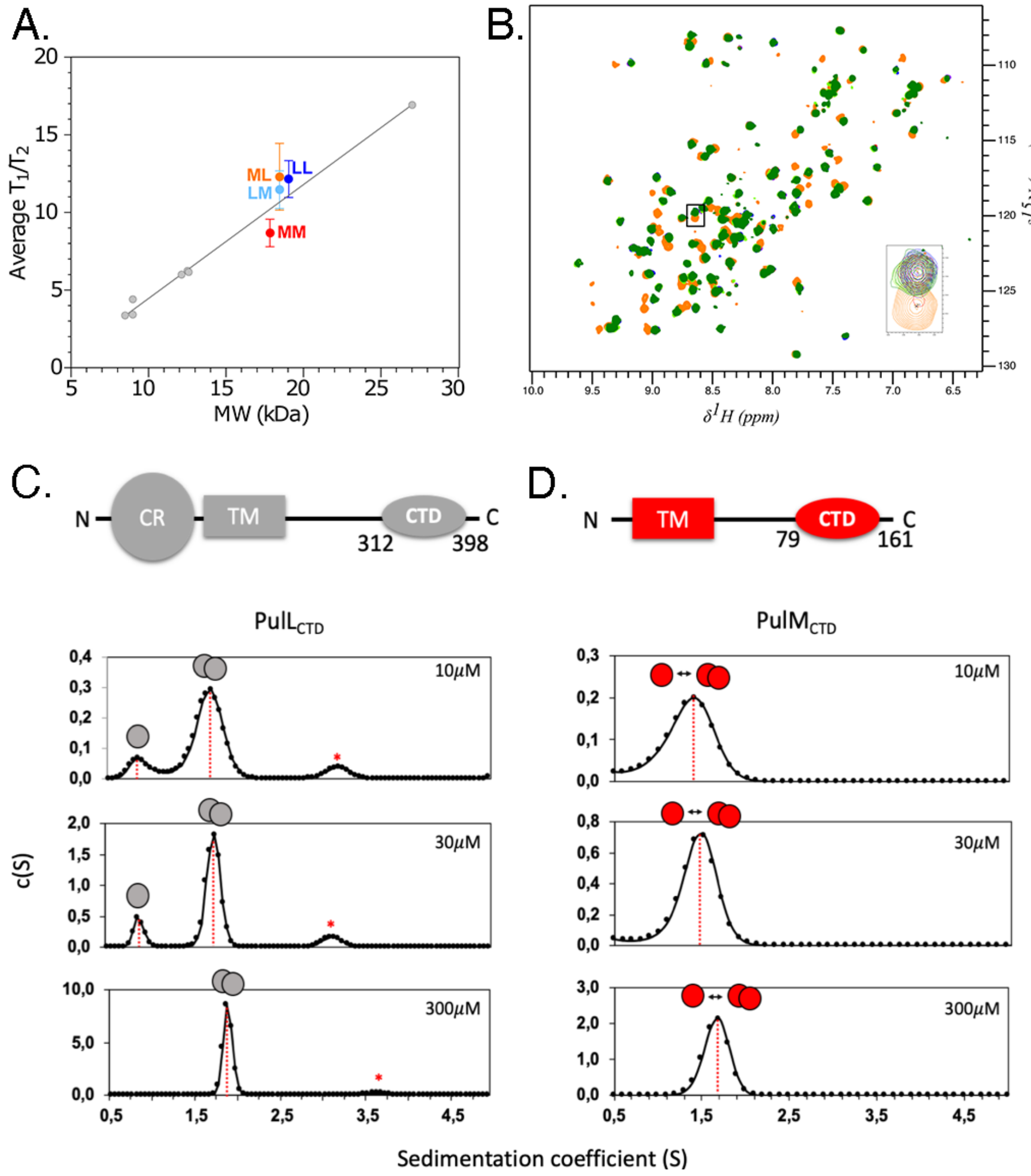


**A.** Average $T_1/T_2$ relaxation times ratio plotted as a function of the molecular weight (MW) are presented for the dimeric states of $PulM_{CTD}$ (red, MM) and $PulL_{CTD}$ (dark blue, LL). For the mixture $PulM_{CTD}$-$PulL_{CTD}$ (orange, ML) and $PulL_{CTD}$-$PulM_{CTD}$ (light blue, LM) the measurements were performed on $^{15}N$-labeled $PulM_{CTD}$ (with unlabeled $PulL_{CTD}$) or on $^{15}N$-labeled $PulL_{CTD}$ (with unlabeled $PulM_{CTD}$), respectively (see Material and Methods). For comparison, grey circles highlight data recorded under similar conditions, at 25°C and 600 MHz, on other in-house protein samples such as human ubiquitin (8.5 kDa, PDB id 1UBQ);

*Klebsiella oxytoca* PulG (12.1 kDa PDB id 5O2Y), etc. **B.** $PulL_{CTD}$-$PulM_{CTD}$ titration by NMR. Left: Superimposition of $^{1}H$-$^{15}N$ HSQC of $^{15}N$-$PulL_{CTD}$ at 100μM in 50mM HEPES pH6.5, 50mM NaCl (orange), in the presence of $PulM_{CTD}$ at 50μM (red), 100μM (purple), 150μM (blue), 200μM $PulM_{CTD}$ (dark green), and 300μM $PulM_{CTD}$ (light green). All spectra were acquired on a 600Mz spectrometer with same parameters. The insert shows a zoom of a representative signal (Asp 331). C. **Determination of $PulL_{CTD}$ oligomeric state by analytical ultracentrifugation (AUC).** Top: Schematic representation of PulL with the cytoplasmic region (CR), transmembrane segment (TM) and the C-terminal domain (CTD). Bottom: Sedimentation coefficient distribution of $PulL_{CTD}$ obtained by AUC at the indicated concentrations. Red dashed lines highlight the main oligomeric forms, schematized with single (monomer) or double (dimer) grey circles. The red star indicates an additional minor peak corresponding to a higher oligomeric state. **D. Determination of $PulM_{CTD}$ oligomeric state by analytical ultracentrifugation (AUC).** Top: Schematic representation of PulM with the transmembrane segment (TM) and the C-terminal domain (CTD). Bottom: Sedimentation coefficient distribution of $PulM_{CTD}$ forms obtained by AUC at the indicated concentrations. Red dashed lines highlight the main oligomeric forms. The different forms are schematized with single (monomer) or double (dimer) red circles.

**Supplementary Figure S2:** Superposition of PulM$_{CTD}$ X-ray structure with PulM$_{CTD}$ NMR structure

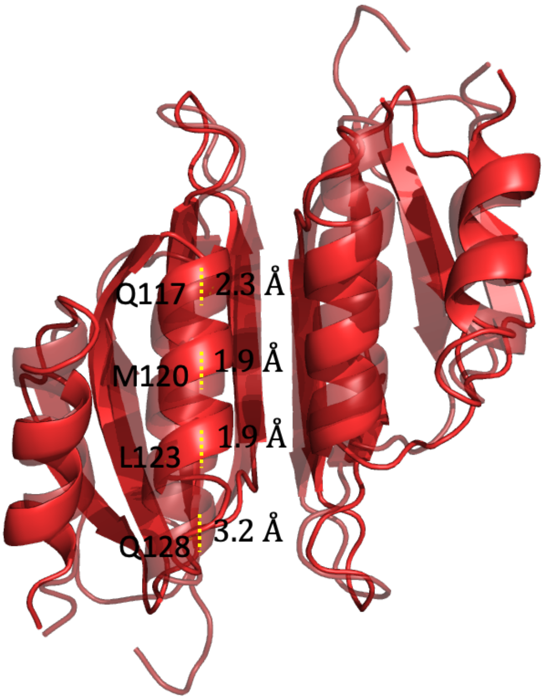


Superposition of PulM$_{CTD}$ X-ray structure (opaque red) with PulM$_{CTD}$ NMR structure (transparent red). The average backbone RMSD is 1.6 Å when the secondary structure elements of both chains are considered. The most noticeable difference between the NMR and X-ray crystallographic structures is a slight displacement of the helix $\alpha$2, 2.3 Å on average along the helix axis. In dashed yellow line, measurement of the position of C$\alpha$ of the helix $\alpha$2 residues between the X-ray structure and the NMR structure of PulM$_{CTD}$.

**Supplementary Figure S3:** Estimation of the PulL and PulM molar ratio in the bacteria.

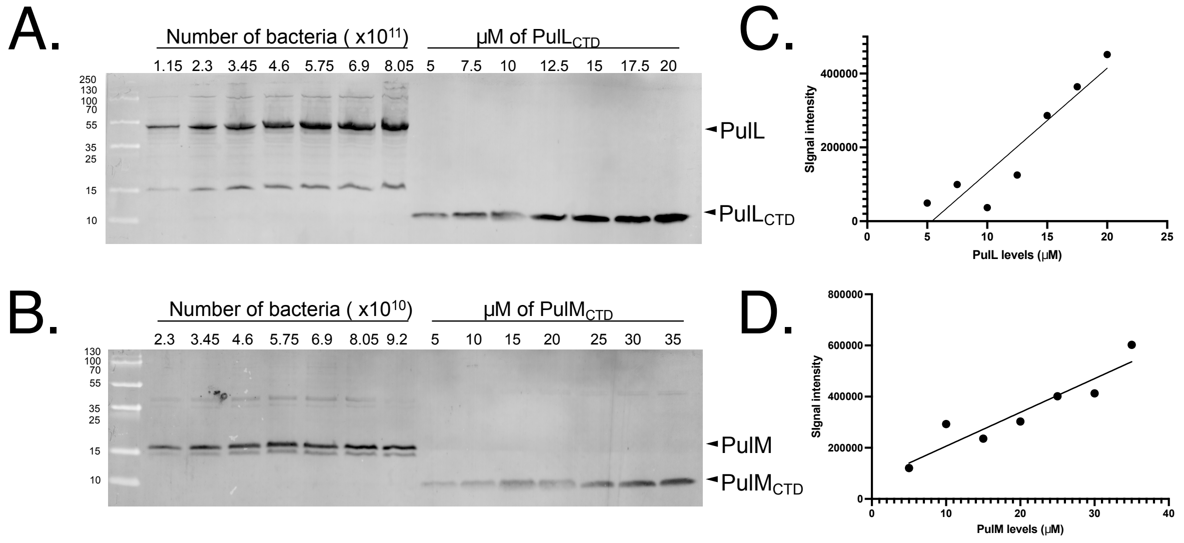


Bacteria of strain PAP7460 carrying plasmid pCHAP8185 were grown to early exponential phase under conditions inducing the *pul* gene expression. Colony forming units were enumerated and total bacterial extracts were prepared from the same cultures at concentrations of 1 and 10 $OD_{600nm}$.m. $L^{-1}$. **A.** Western blots of total bacterial extracts from the numbers of bacteria indicated above each lane and known concentrations of purified $PulL_{CTD}$, revealed with anti-PulL antibodies. Positions of PulL and $PulL_{CTD}$ are indicated on the right. The Mw standards are shown on the left (in kDa). **B.** Western blots of total bacterial extracts from the number of bacteria indicated above each lane and known concentrations of purified $PulM_{CTD}$, revealed with anti-PulM antibodies. **C**. Standard curve quantifying the signal intensity of known amounts of purified $PulL_{CTD}$ (in µM) analyzed in panel (**A**). **D**. Standard curve quantifying the signal intensity of known amounts of purified $PulM_{CTD}$ (in µM) analyzed in panel (**B**).

**Supplementary Figure S4:** Homologues and orthologues of PulL

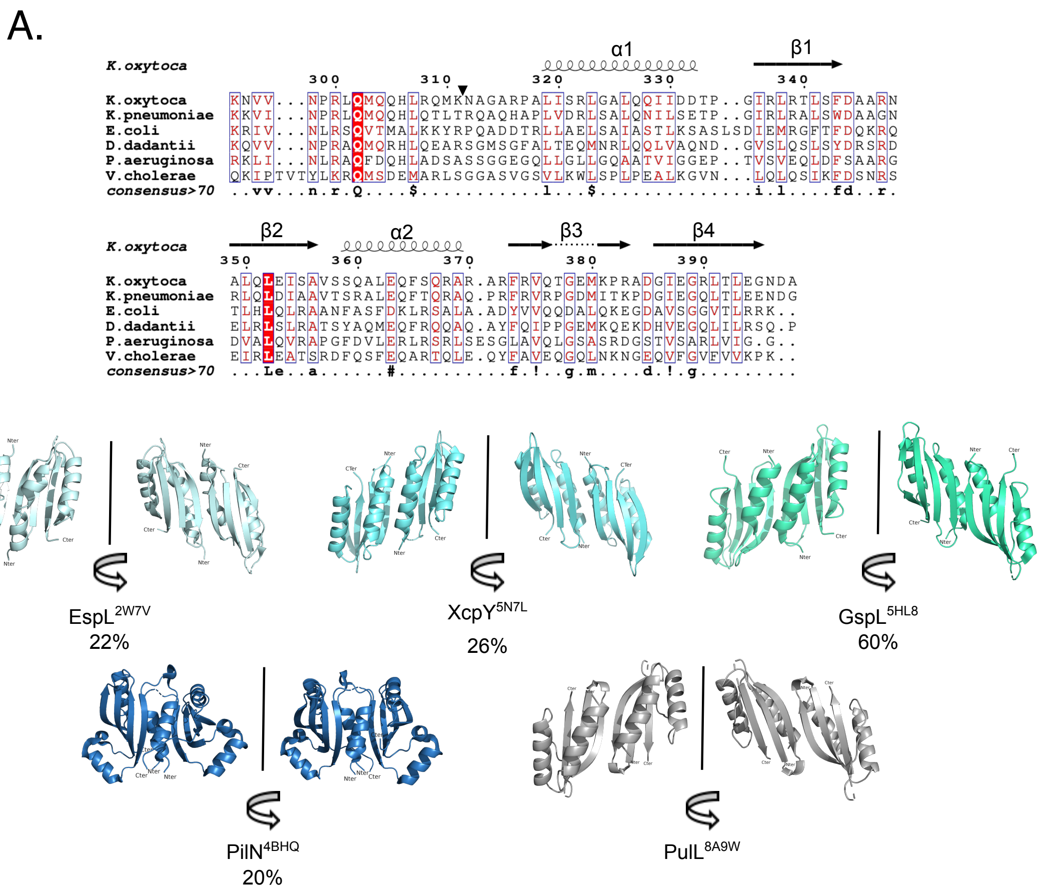


**A**. Sequence alignment of representative examples of the C-terminal domain of PulL homologues and orthologues. The residue numbering is for PulL, and the triangle indicates the beginning of the C-terminal domain of PulL used in this study. The secondary structure elements correspond to the NMR structure of the PulL. The alignment was performed by using Escript (Robert and Gouet, 2014). **B**. Structures of the $PulL_{CTD}$ homologues and orthologues obtained by X-ray crystallography. The percentage of identity with $PulL_{CTD}$ is indicated below each structure.

**Supplementary Figure S5:** Homologues and orthologues of PulM

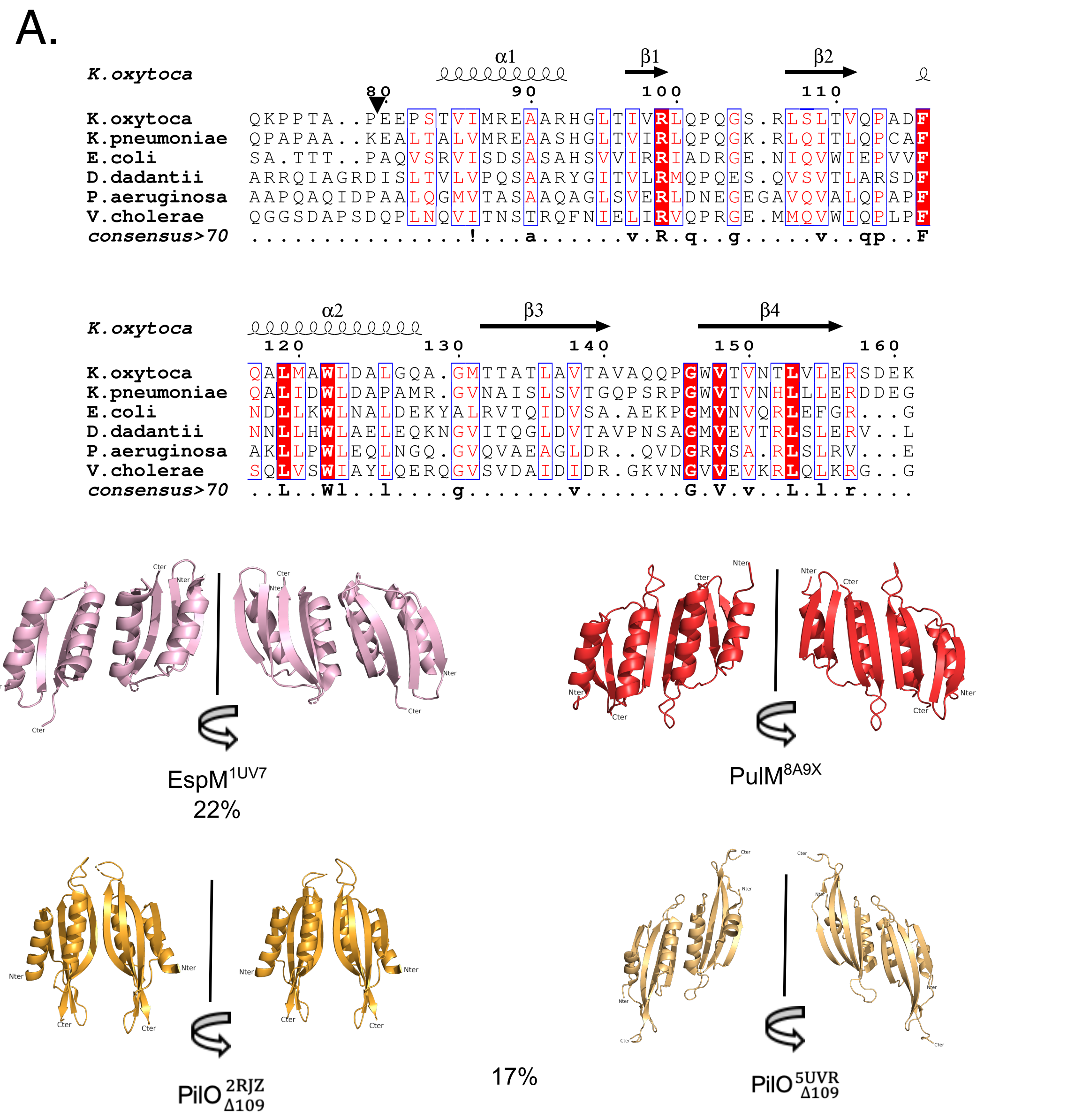


**A.** Sequence alignment of representative examples of the C-terminal domain of PulM homologues and orthologues. The residue numbering is for PulM; the triangle corresponds to the beginning of the CTD of PulM used in this study. The secondary structure elements correspond to the NMR structure of PulM$_{CTD}$. The alignment was performed by using Escript (Robert and Gouet, 2014). **B.** Structures of PulM$_{CTD}$ homologues and orthologues determined by X-ray crystallography. The percentage of identity with PulM$_{CTD}$ is indicated below each structure.

**Supplementary Figure S6:** Structural alignments of $PulM_{CTD}$ and $PulL_{CTD}$ dimer interfaces.

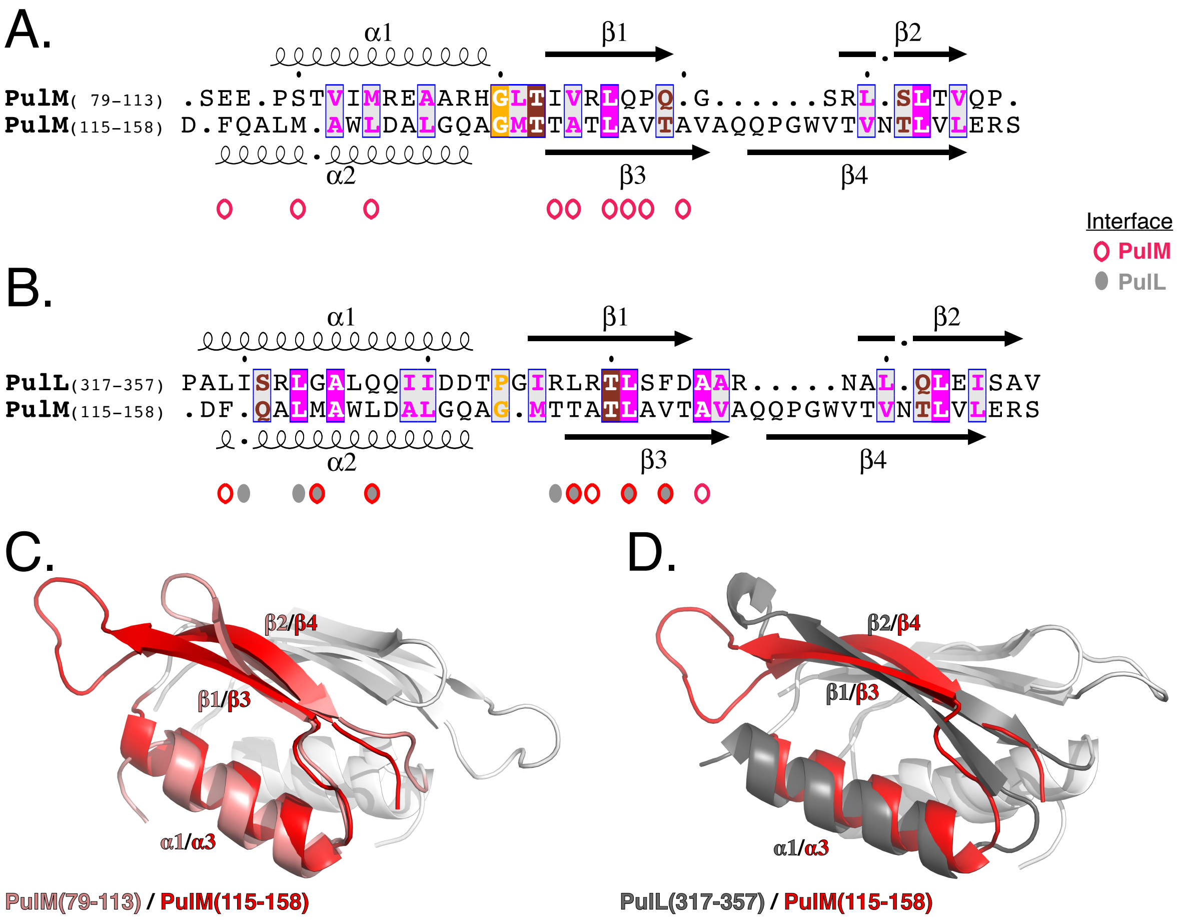


**A.** Sequence alignment based on structural superimposition of $PulM_{CTD}$ (79-113) and $PulM_{CTD}$ (115-158) with TM-align (Zhang and Skolnick, 2005). Position with similar amino acids are boxed and colored by amino acid properties (hydrophobics in pink, polar in maroon, glycine and proline in orange). Secondary structure elements are labelled on top for $PulM_{CTD}$ (79-113) and bottom for $PulM_{CTD}$ (115-158). Positions involved in $PulM_{CTD}$ homodimerization are highlighted with red open circles. **B.** Structural alignment of $PulL_{CTD}$ (317-358) and $PulM_{CTD}$ (115-158), with amnio acids colored as in (A). Secondary structure elements are labelled on top for $PulL_{CTD}$ (317-358) and bottom for $PulM_{CTD}$ (115-158). Positions involved in $PulL_{CTD}$-$PulM_{CTD}$ heterodimerization are highlighted with filled grey circles for PulL and red open circles for PulM. **C.** Superimposition of $PulM_{CTD}$ (79-113) in pink and $PulM_{CTD}$ (115-158) in red. **D.** Superimposition of $PulL_{CTD}$ (317-357) in grey and $PulM_{CTD}$ (115-158) in red. In C and D, secondary structure elements are labeled in the respective colors. Parts of the proteins not used for superimposition are colored in white for sake of clarity.

**Supplementary Figure S7:** Levels of PulL and PulM variants in bacterial extracts.

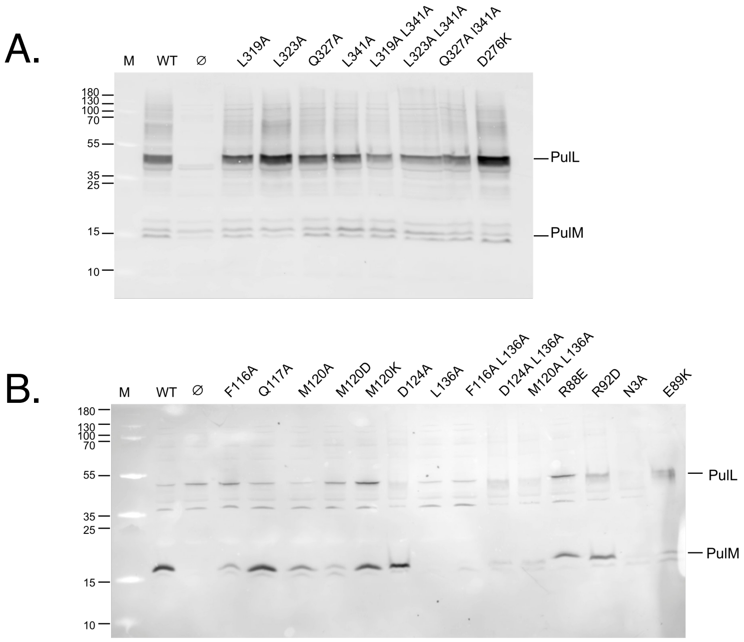


**A.** Total extracts of strain PAP7460 harboring pCHAP8251 (lacking the *pulL* gene) complemented with plasmids encoding PulL or its mutant variants. Total extracts from 0.1 $OD_{600nm}$ of cultures were analyzed on 10% Tris-Tricine SDS-PAGE followed by Western blot with anti-PulM and anti-PulL antibodies. **B.** Total extracts of strain PAP7460 harboring pCHAP8496 (lacking the *pulM* gene) complemented with plasmids encoding PulM or its mutant variants. Total extracts from 0.1 $OD_{600nm}$ of cultures were analyzed on 10% Tris-Tricine SDS-PAGE followed by Western blot with anti-PulM and anti-PulL antibodies. Note that the L136A substitution in PulM affects the epitope recognized by anti-PulM antibodies.

**Supplementary Figure S8:** Sequence and structure analysis used for PulL-PulM complex modeling.

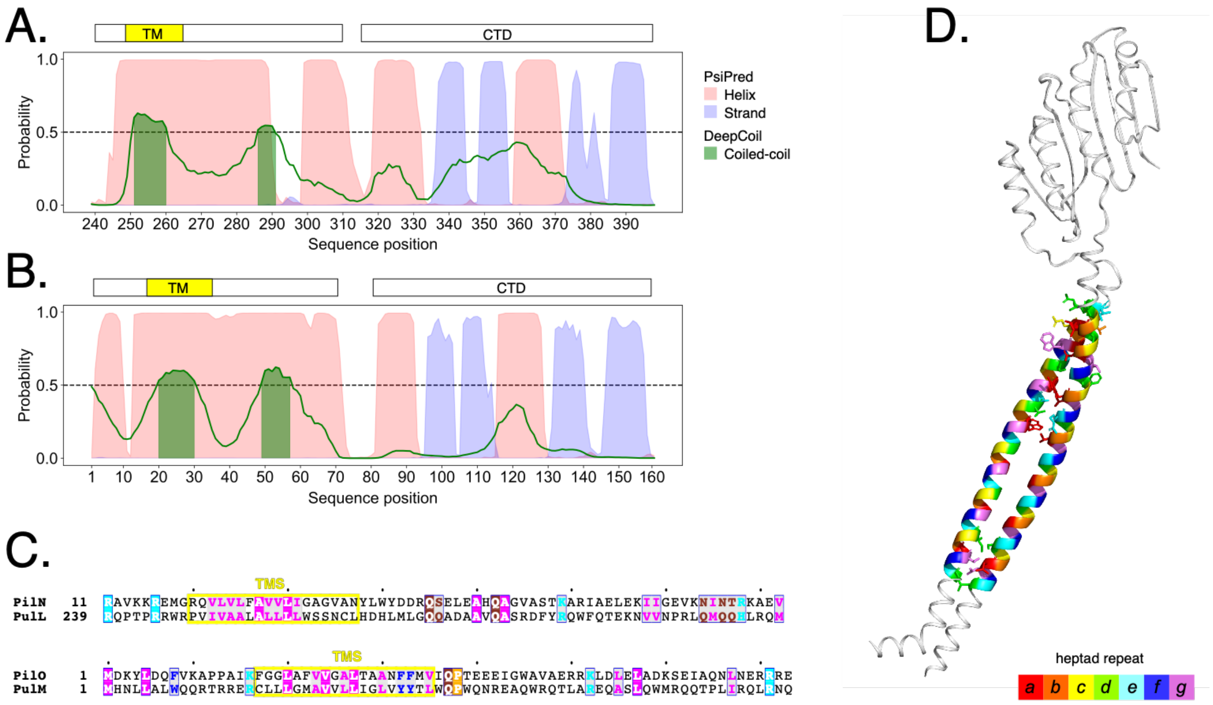


Probabilities for secondary structures [PsiPred] (Jones, 1999) and coiled-coil [DeepCoil] (Ludwiczak et al., 2019) predictions for PulL (**A**) and PulM (**B**) sequences. Helical probabilities are shown in red, beta-strand in blue and coiled-coil in green. Position of the transmembrane segments (TM) is shown in yellow. Regions with coiled-coil probability above 0.5 are highlighted in green. **C.** Sequence alignments of the N-terminal regions of *M. xanthus* $PilN_{11-81}$/$PilO_{1-73}$ with $PulL_{239-310}$/$PulM_{1-73}$ as used for modeling. Positions with similar amino acids are boxed and colored by amino acid properties: hydrophobic in pink, polar in maroon, positively charged in cyan, aromatics in blue, and glycine and proline in orange. The predicted TMS are highlighted in yellow. **D.** Knobs-into-Holes and heptad repeats found in the model of the PulL-PulM complex using the SOCKET2 program (Kumar and Woolfson, 2021). Residues are colored according to their positions in the heptad repeat [a-g] of the coiled-coil. Side-chains of residues detected as knobs or forming holes are shown as sticks.

**Supplementary Table S1**: PulL$_{CTD}$ monomer NMR structure statistics and restraints.

| | |
|---|---|
| **Number of restraints** | |
| NOE Distance restraints | |
| Intra-residue (\|i-j\| = 0) | 392 |
| Sequential (\|i-j\| = 1) | 262 |
| Medium-range (2 ≤ \|i-j\| < 5) | 116 |
| Long-range (\|i-j\| ≥ 5) | 170 |
| Ambiguous | 23 |
| *Total* | 963 |
| Dihedral angle restraints (φ/ψ) | 144 (72/72) |
| Hydrogen bonds restraints | 28 |
| **Restraints statistics[a]** | |
| Average no. of violations per structure | |
| NOE restraints >0.5 Å | 0.7 ± 0.8 |
| H-bond restraints >0.5 Å | 0 |
| Dihedral restraints >5 ° | 1.1 ± 1.4 |
| RMS of distance violations | |
| NOE restraints | 0.053 ± 0.013 Å |
| H-bond restraints | 0.034 ± 0.004 Å |
| RMS of dihedral violations | 1.009 ± 0.408 ° |
| **RMS from idealized covalent geometry** | |
| bonds | 0.005 ± 0.001 Å |
| angles | 0.582 ± 0.035 ° |
| impropers | 1.639 ± 0.150 ° |
| **Structural quality[a]** | |
| *Ramachandran statistics[b]* | |
| Most favoured regions | 89.4 ± 2.8 % |
| Allowed regions | 10.1 ± 3.2 % |
| Generously allowed regions | 0.4 ± 0.6 % |
| Disallowed regions | 0.1 ± 0.4 % |
| *Global quality scores (Raw / Z-score)[c]* | |
| Verify3D | 0.20 / -4.17 |
| ProsaII | 0.52 / -0.54 |
| ProCheck (all) | -0.44 / -2.60 |
| MolProbity clashscore | 38.6 / -5.10 |
| **Coordinates precision[d]** | |
| Backbone atoms (311-398) | 1.44 ± 0.40 Å |
| Heavy atoms (311-398) | 1.74 ± 0.38 Å |
| Backbone atoms (317-344, 346-385,3 87-395[e]) | 0.60 ± 0.21 Å |
| Heavy atoms (317-344, 346-385, 387-395[e]) | 1.13 ± 0.30 Å |

[a] Average values and standard deviations over the 10 conformers
[b] Percentage of residues in the Ramachandran plot regions determined by PROCHECK (Laskowski et al., 1996)
[c] Calculated using PSVS ver. 1.5 (Bhattacharya et al., 2007)
[d] Average root mean square deviation (RMSD) over the 10 conformers' atomic coordinates with respect to the average structure.
[e] Ordered residues [S(phi) + S(psi) > 1.8].

**Supplementary Table S2:** Data collection and refinement statistics.

| Parameters | PulL$_{CTD}$ | PulL$_{CTD}$-PulM$_{CTD}$ | PulM$_{CTD}$ |
|---|---|---|---|
| **Beamline** | Proxima 2A | Proxima 1 | Proxima 1 |
| **Resolution range** | 40.15 - 1.895 (1.963 - 1.895) | 58.95 - 2.771 (2.87 - 2.771) | 40.4 - 1.52 (1.575 - 1.52) |
| **Space group** | I 21 3 | P 63 2 2 | C 2 2 21 |
| **Unit cell (Å, °)** | 80.3 80.3 80.3 90 90 90 | 117.899 117.899 110.002 90 90 120 | 80.81 135.48 109.4 90 90 90 |
| **Total reflections** | 277247 (26424) | 464974 (47811) | 1239705 (112086) |
| **Unique reflections** | 7012 (687) | 11980 (1165) | 91879 (8852) |
| **Multiplicity** | 39.5 (38.5) | 38.8 (41.0) | 13.5 (12.7) |
| **Completeness (%)** | 99.97 (100.00) | 99.92 (100.00) | 99.60 (96.68) |
| **Mean I/sigma(I)** | 33.73 (3.03) | 29.12 (2.73) | 18.95 (1.52) |
| **Wilson B-factor** | 41.28 | 95.24 | 27.69 |
| **R-merge** | 0.06935 (1.739) | 0.09584 (1.471) | 0.06888 (1.059) |
| **R-meas** | 0.07029 (1.762) | 0.09716 (1.49) | 0.07162 (1.102) |
| **R-pim** | 0.01132 (0.2832) | 0.01573 (0.2311) | 0.01943 (0.3013) |
| **CC1/2** | 1 (0.86) | 1 (0.846) | 0.999 (0.837) |
| **CC*** | 1 (0.962) | 1 (0.958) | 1 (0.955) |
| **Reflections used in refinement** | 7012 (687) | 11980 (1165) | 91879 (8830) |
| **Reflections used for R-free** | 339 (32) | 581 (50) | 4593 (442) |
| **R-work** | 0.2408 (0.3641) | 0.2569 (0.3558) | 0.2082 (0.4667) |

| | | | |
|---|---|---|---|
| **R-free** | 0.2765 (0.2755) | 0.2980 (0.4413) | 0.2321 (0.4923) |
| **CC(work)** | 0.946 (0.725) | 0.907 (0.681) | 0.955 (0.801) |
| **CC(free)** | 0.915 (0.713) | 0.952 (0.552) | 0.949 (0.779) |
| **Number of non-hydrogen atoms** | 606 | 2365 | 4696 |
| **Macromolecules** | 579 | 2326 | 4141 |
| **Ligands** | 10 | 0 | 0 |
| **Solvent** | 17 | 39 | 555 |
| **Protein residues** | 75 | 308 | 556 |
| **RMS(bonds)** | 0.010 | 0.012 | 0.024 |
| **RMS (angles)** | 1.41 | 1.62 | 3.61 |
| **Ramachandran favored (%)** | 94.37 | 90.00 | 95.74 |
| **Ramachandran allowed (%)** | 2.82 | 8.00 | 2.78 |
| **Ramachandran outliers (%)** | 2.82 | 2.00 | 1.48 |
| **Rotamer outliers (%)** | 10.00 | 10.29 | 2.07 |
| **Clashscore** | 11.83 | 9.52 | 14.79 |
| **Average B-factor** | 64.98 | 122.89 | 35.56 |
| **Macromolecules** | 65.09 | 123.41 | 35.13 |
| **Ligands** | 79.76 | | |
| **Solvent** | 52.56 | 92.02 | 38.74 |
| **Number of TLS groups** | | | 39 |

Statistics for the highest-resolution shell are shown in parentheses.

**Supplementary Table S3:** PulM$_{CTD}$ dimer NMR structure statistics and restraints.

| | |
|---|---|
| **Number of restraints (per monomer)** | |
| NOE Distance restraints | |
| Intra-residue (\|i-j\| = 0) | 501 |
| Sequential (\|i-j\| = 1) | 324 |
| Medium-range (2 ≤ \|i-j\| < 5) | 184 |
| Long-range (\|i-j\| ≥ 5) | 455 |
| Inter-molecular | 25 |
| Ambiguous | 159 |
| *Total* | 1648 |
| Dihedral angle restraints (φ/ψ) | 138 (69/69) |
| Hydrogen bonds restraints | 12 |
| **Restraints statistics[a]** | |
| Average no. of violations per structure | |
| NOE restraints >0.5 Å | 0 |
| H-bond restraints >0.5 Å | 0 |
| Dihedral restraints >5 ° | 0 |
| RMS of distance violations | |
| NOE restraints | 0.046 ± 0.001 Å |
| H-bond restraints | 0.021 ± 0.001 Å |
| RMS of dihedral violations | 0.869 ± 0.083 ° |
| **RMS from idealized covalent geometry** | |
| bonds | 0.007 ± 0.001 Å |
| angles | 0.793 ± 0.009 ° |
| impropers | 1.948 ± 0.039 ° |
| **Structural quality[a]** | |
| *Ramachandran statistics[b]* | |
| Most favoured regions | 93.5 ± 1.3 % |
| Allowed regions | 5.9 ± 1.2 % |
| Generously allowed regions | 0 ± 0 % |
| Disallowed regions | 0.6 ± 0.7 % |
| *Global quality scores (Raw / Z-score)[c]* | |
| Verify3D | 0.15 / -4.98 |
| ProsaII | 0.85 / 0.83 |
| ProCheck (all) | -0.34 / -2.01 |
| MolProbity clashscore | 48.8 / -6.85 |
| **Coordinates precision[d]** | |
| Backbone atoms (79-161) | 0.69 ± 0.14 Å |
| Heavy atoms (79-161) | 1.10 ± 0.16 Å |
| Backbone atoms (84-101,106-160[e]) | 0.35 ± 0.09 Å |
| Heavy atoms (84-101,106-160[e]) | 0.69 ± 0.07 Å |

[a] Average values and standard deviations over the 15 conformers
[b] Percentage of residues in the Ramachandran plot regions determined by PROCHECK (Laskowski et al., 1996)
[c] Calculated using PSVS ver. 1.5 (Bhattacharya et al., 2007)
[d] Average root mean square deviation (RMSD) over the 15 conformers' atomic coordinates with respect to the average structure.
[e] Ordered residues [S(phi) + S(psi) > 1.8].

**Supplementary Table S4:** Scoring biological interfaces in the X-ray crystallographic structure of the $\text{PulL}_{\text{CTD}}$-$\text{PulM}_{\text{CTD}}$ heterodimer complex

| | $\mathbf{PulL}^{3}_{CTD}$-$\mathbf{PulM}^{4}_{\mathbf{CTD}}$ | $\mathbf{PulL}^{3}_{CTD}$-$\mathbf{PulM}^{2}_{\mathbf{CTD}}$ |
|---|---|---|
| **Prodigy-Crystal** | 19% | 7% |
| **ClusPro-DC** | 58% | 11% |

We tested the probability of biological relevance of heterodimers formed by $\text{PulL}^{3}_{CTD}$ and molecules $\text{PulM}^{4}_{\text{CTD}}$ or $\text{PulM}^{2}_{\text{CTD}}$ within the asymmetric unit. Both Prodigy-Crystal ((Jiménez-García et al., 2019) and ClusPro-DC (Yueh et al., 2017) predicted a higher probability for $\text{PulL}^{3}_{CTD}$-$\text{PulM}^{4}_{\text{CTD}}$ to be a biologically relevant interface than $\text{PulL}^{3}_{CTD}$-$\text{PulM}^{2}_{\text{CTD}}$. These results agree with PISA analysis of the buried interface for each heterodimer and, more importantly, with the results of the NMR CSP analysis of heterodimer interface in solution.

**Supplementary Table S5:** Plasmids used in this study

| Plasmid | Markers | Source/Reference |
|---|---|---|
| pSU18 | p*lacZ*, p15A *ori*, Cm$^R$ | (Bartolomé et al., 1991) |
| pMalP2 | p*lacZ*, *malE*, ColE1 *ori*, Ap$^R$ | New England Biolabs |
| pPulL$_{CTD}$ | pMalP2-His-TEV-PulL$_{CTD}$ | (Dazzoni et al., 2021) |
| pPulM$_{CTD}$ | pMalP2-His-TEV-PulM$_{CTD}$ | *Proteogenix* |
| pCHAP8185 | *pulS ,pulAB*, *pulCDEFGHIJKLMNO,* ColE1 *ori,* Ap$^R$ | (Cisneros et al., 2012) |
| pCHAP8496 | pCHAP8185 *ΔpulM*, Ap$^R$ | (Santos-Moreno et al., 2017) |
| pCHAP8251 | pCHAP8185 *ΔpulL*, Ap$^R$ | (Santos-Moreno et al., 2017) |
| pCHAP8258 | pSU18-*pulL,* Cm$^R$ | (Santos-Moreno et al., 2017) |
| pCHAP1353 | pSU18-*pulM*, Cm$^R$ | (Possot et al., 2000) |
| pMS1185 | pSU18-*pulL* L319A, Cm$^R$ | This study |
| pMS1186 | pSU18-*pulL* L323A, Cm$^R$ | This study |
| pMS1187 | pSU18-*pulL* Q327A, Cm$^R$ | This study |
| pMS1188 | pSU18-*pulL* L341A, Cm$^R$ | This study |
| pMS1235 | pSU18-*pulL* L319A L341A, Cm$^R$ | This study |
| pMS1236 | pSU18-*pulL* L323A L341A, Cm$^R$ | This study |
| pMS1237 | pSU18-*pulL* Q327A L341A, Cm$^R$ | This study |
| pMS1239 | pSU18-*pulL* Q327A L319A, Cm$^R$ | This study |
| pMS1189 | pSU18-*pulM* F116A, Cm$^R$ | This study |
| pMS1190 | pSU18-*pulM* M120A, Cm$^R$ | This study |
| pMS1166 | pSU18-*pulM* M120D, Cm$^R$ | This study |
| pCHAP8705 | pSU18-*pulM* M120K, Cm$^R$ | This study |
| pMS1192 | pSU18-*pulM* D124A, Cm$^R$ | This study |
| pMS1201 | pSU18-*pulM* L136A, Cm$^R$ | This study |
| pMS1223 | pSU18-*pulM* D124A L136A, Cm$^R$ | This study |
| pMS1224 | pSU18-*pulM* F116A L136A, Cm$^R$ | This study |
| pMS1225 | pSU18-*pulM* M120A L136A, Cm$^R$ | This study |
| pCHAP8468 | pSU18-*pulM* R88E, Cm$^R$ | This study |
| pCHAP8525 | pSU18-*pulM* R92D, Cm$^R$ | This study |
| pCHAP8728 | pSU18-*pulM* N3A, Cm$^R$ | This study |
| pMS1297 | pSU18 *pulL* C264L Cm$^R$ | This study |
| pMS1296 | pSU18 *pulL* 249C C264L, Cm$^R$ | This study |
| pMS1290 | pSU18 *pulL* 250C C264L, Cm$^R$ | This study |
| pMS1291 | pSU18 *pulL* 251C C264L, Cm$^R$ | This study |
| pMS1292 | pSU18 *pulL* 252C C264L, Cm$^R$ | This study |
| pMS1293 | pSU18 *pulL* 253C C264L, Cm$^R$ | This study |
| pMS1294 | pSU18 *pulL* 254C C264L, Cm$^R$ | This study |
| pMS1295 | pSU18 *pulL* 255C C264L, Cm$^R$ | This study |
| pMS1300 | pSU18 *pulL* 256C C264L, Cm$^R$ | This study |
| pCHAP8813 | pUC18 *pulM* (17C), Ap$^R$ | This study |
| pCHAP8893 | pUC18 *pulM* C17L, Ap$^R$ | This study |
| pCHAP8894 | pUC18 *pulM* C17L L18C, Ap$^R$ | This study |
| pCHAP8817 | pUC18 *pulM* C17L L19C, Ap$^R$ | This study |
| pCHAP8818 | pUC18 *pulM* C17L L20C, Ap$^R$ | This study |

| | | |
|---|---|---|
| pCHAP8819 | pUC18 *pulM* C17L G21C, Ap$^R$ | This study |
| pCHAP8820 | pUC18 *pulM* C17L M22C, Ap$^R$ | This study |
| pCHAP8821 | pUC18 *pulM* C17L A23C, Ap$^R$ | This study |
| pCHAP8822 | pUC18 *pulM* C17L V24C, Ap$^R$ | This study |
| pUT18C | T18 *cyaA* fragment, ColE1 *ori*, Ap$^R$ | (Karimova G et al., 1998) |
| pKT25 | T25 *cyaA* fragment, p15A *ori,* Km$^R$ | (Karimova G et al., 1998) |
| pUT18C-Zip | pUT18C fused to yeast Leu zipper motif, Ap$^R$ | (Karimova G et al., 1998) |
| pKT25-Zip | pKT25 with yeast Leu zipper Km$^R$ | (Karimova G et al., 1998) |
| pMS1222 | pUT18C-*pulL*, Ap$^R$ | This study |
| pMS1229 | pKT25-*pulL*, Km$^R$ | This study |
| pCHAP8154 | pUT18C-*pulM,* Ap$^R$ | (Nivaskumar et al., 2016) |
| pCHAP8155 | pKT25-*pulM,* Km$^R$ | (Nivaskumar et al., 2016) |

**Supplementary Table S6:** Oligonucleotides used in this study

| Oligonucleotide name | DNA sequence (5'-3') |
|---|---|
| PulL Kpn 5 | GCAGGTACCTATGAATAACCACCATACC |
| PulL Eco 3 | CACGAATTCCTGTTGCCATAAGGCGAG |
| PulL L319A For* | GGCGCCCGGCCGGCGGCTATCTCGCGGCTCGGCGC |
| PulL L323A For | CGCTTATCTCGCGGGCCGGCGCGCTGCAGCAAATC |
| PulL Q327A For | CGGCTCGGCGCGCTGGCGCAAATCATCGACGACAC |
| PulL L341A For | CATCCGCCTGCGGACGGCGAGCTTTGACGCCGCGC |
| PulM F116A For | GTGCAGCCCGCCGATGCCCAGGCGTTGATGGCATG |
| PulM M120A For | CGATTTCCAGGCGTTGGCGGCATGGCTGGACGCGC |
| PulM M120D For | GATTTCCAGGCGTTGGACGCATGGCTGGACGCG |
| PulM M120K For | CGATTTCCAGGCGTTGAAGGCATGGCTGGAC |
| PulM D124A For | CGTTGATGGCATGGCTGGCCGCGCTGGGGCAGGC |
| PulM L136A For | GATGACCACCGCCACCGCGGCGGTGACCGCCGTAG |
| PulM R88E For | GACGGTGATTATGGACGAAGCCGCCCGTC |
| PulM R92D For | CGCGAAGCCGCCGATCACGGTCTCACCATCG |
| PulM N3A For | GGAACGATGCATGCCCTGCTCGCCTTATG |
| PulL C264L For | CTGTGGAGCAGTAATCTCC TCCACGATCATC |
| PulL V249C For | CGACGCTGGCGCCCGTGTATCGTCGCGGCGCTG |
| PulL I250C For | CTGGCGCCCGGTATGCGTCGCGGCGCTG |
| PulL V251C For | CTGGCGCCCGGTAATCTGCGCGGCGCTGGCGCTAC |
| PulL A252C For | CGCCCGGTAATCGTCTGCGCGCTGGCGCTAC |
| PulL A253C For | CGGTAATCGTCGCGTGCCTGGCGCTACTGCTGC |
| PulL L254C For | GTAATCGTCGCGGCGTGTGCGCTACTGCTGC |
| PulL A255C For | CGTCGCGGCGCTGTGCCTACTGCTGCTGTG |
| PulL L256C For | GTCGCGGCGCTGGCGTGTCTGCTGCTGTGGAG |
| PulM C17L For | GCCGCGAACGGCTTCTGCTGCTGG |
| PulM C17L L18C For | CCGCGAACGGCTTTGTCTGCTGGGTATGG |
| PulM C17L L19C For | CCGCGAACGGCTTCTGTGTCTGGGTATGGCC |
| PulM C17L L20C For | CGAACGGCTTCTGCTGTGTGGTATGGCCGTGG |
| PulM C17L G21C For | CGGCTTCTGCTGCTGTGTATGGCCGTGGTAC |
| PulM C17L M22C For | CTTCTGCTGCTGGGTTGTGCCGTGGTACTGC |
| PulM C17L A23C For | GCTGCTGGGTATGTGCGTGGTACTGCTC |
| PulM C17L V24C For | CTGCTGGGTATGGCCTGTGTACTGCTCATCG |

* Only the Forward primers used for site-directed mutagenesis are shown. They were used in combination with Reverse primers with a fully overlapping antiparallel sequence.

### Supplementary Materials and Methods

#### NMR relaxation experiments

The $^{15}N$ relaxation times ($T_1$ and $T_2$) were estimated at 25°C on the 600 MHz spectrometer on samples of $PulL_{CTD}$ (120 µM of $^{15}N$-$PulL_{CTD}$, pH 6.5), $PulM_{CTD}$ (30 µM of $^{15}N$-$PulM_{CTD}$, pH 7.0), $PulL_{CTD}$-$PulM_{CTD}$ complex (30 µM of $^{15}N$-$PulL_{CTD}$ and 120 µM of unlabeled-$PulM_{CTD}$, pH 6.5) and $PulM_{CTD}$-$PulL_{CTD}$ complex (30 µM of $^{15}N$-$PulM_{CTD}$ and 120 µM of unlabeled-$PulL_{CTD}$, pH 7.0). Experiments were recorded by standard methods (Barbato *et al.*, 1992) in an interleaved manner with a recycling time of 4.5 s. Two relaxation delays for $T_1$ (20 and 500 ms) and $T_2$ (17 and 51 ms) were used with 184 scans. For each residue, the $T_1$ and $T_2$ were estimated from the peak intensities ($Int_1$ and $Int_2$) for the two delays ($d_{relax1}$ and $d_{relax2}$) by the formula $(d_{relax1} - d_{relax2})/\ln(Int_2/Int_1)$. The overall $T_1$ and $T_2$ values (and errors) for each sample were calculated as the averages (and standard deviations) over all non-flexible residues.

**Analytical ultracentrifugation experiments**

Sedimentation velocity experiments were carried out at 20°C using a Beckman Coulter Optima AUC centrifuge equipped with an AN60-Ti rotor. Protein samples at different concentrations (10 µM, 30 µM, 300 µM) were loaded in a 1.2 cm or 3 mm centerpiece and centrifuged overnight at 42000 rpm. Data were analyzed with SEDFIT 15.1 using a continuous size distribution c(S) model. The partial specific volume, the viscosity and the density of the samples were calculated with SEDNTERP software. The processed data were used to obtain sedimentation coefficients values at null concentration in our experimental conditions ($S_0$), and to get the standard sedimentation coefficients in buffer (50 mM HEPES pH 7.0, 50 mM NaCl).

### Supplementary References:


BARBATO, G., IKURA, M., KAY, L., PASTOR, R. & BAX, A. 1992. Backbone dynamics of calmodulin studied by 15N relaxation using inverse detected two-dimensional NMR spectroscopy: the central helix is flexible. *Biochemistry,* 31.

BARTOLOMÉ, B., JUBETE, Y., MARTÍNEZ, E. & DE LA CRUZ, F. 1991. Construction and properties of a family of pACYC184-derived cloning vectors compatible with pBR322 and its derivatives. *Gene,* 102.

BHATTACHARYA, A., TEJERO, R. & MONTELIONE, G. 2007. Evaluating protein structures determined by structural genomics consortia. *Proteins,* 66.

CISNEROS, D., BOND, P., PUGSLEY, A., CAMPOS, M. & FRANCETIC, O. 2012. Minor pseudopilin self-assembly primes type II secretion pseudopilus elongation. *The EMBO journal,* 31.

JIMÉNEZ-GARCÍA, B., ELEZ, K., KOUKOS, P., BONVIN, A. & VANGONE, A. 2019. PRODIGY-crystal: a web-tool for classification of biological interfaces in protein complexes. *Bioinformatics (Oxford, England),* 35.

JONES, D. T. 1999. Protein secondary structure prediction based on position-specific scoring matrices. *J Mol Biol,* 292**,** 195-202.

KARIMOVA G, PIDOUX J, ULLMANN A & LADANT D 1998. A bacterial two-hybrid system based on a reconstituted signal transduction pathway. *Proceedings of the National Academy of Sciences of the United States of America,* 95.

KUMAR, P. & WOOLFSON, D. N. 2021. Socket2: A Program for Locating, Visualising, and Analysing Coiled-coil Interfaces in Protein Structures. *Bioinformatics*.

LASKOWSKI, R., RULLMANNN, J., MACARTHUR, M., KAPTEIN, R. & THORNTON, J. 1996. AQUA and PROCHECK-NMR: programs for checking the quality of protein structures solved by NMR. *Journal of biomolecular NMR,* 8.

LUDWICZAK, J., WINSKI, A., SZCZEPANIAK, K., ALVA, V. & DUNIN-HORKAWICZ, S. 2019. DeepCoil-a fast and accurate prediction of coiled-coil domains in protein sequences. *Bioinformatics,* 35**,** 2790-2795.

NIVASKUMAR, M., SANTOS-MORENO, J., MALOSSE, C., NADEAU, N., CHAMOT-ROOKE, J., TRAN VAN NHIEU, G. & FRANCETIC, O. 2016. Pseudopilin residue E5 is essential for recruitment by the type 2 secretion system assembly platform. *Mol Microbiol,* 101**,** 924-41.

POSSOT, O., VIGNON, G., BOMCHIL, N., EBEL, F. & PUGSLEY, A. 2000. Multiple interactions between pullulanase secreton components involved in stabilization and cytoplasmic membrane association of PulE. *Journal of bacteriology,* 182.

ROBERT, X. & GOUET, P. 2014. Deciphering key features in protein structures with the new ENDscript server. *Nucleic acids research,* 42.

SANTOS-MORENO, J., EAST, A., GUILVOUT, I., NADEAU, N., BOND, P., TRAN VAN, N. G. & FRANCETIC, O. 2017. Polar N-terminal Residues Conserved in Type 2 Secretion Pseudopilins Determine Subunit Targeting and Membrane Extraction Steps during Fibre Assembly. *Journal of molecular biology,* 429.

ZHANG, Y. & SKOLNICK, J. 2005. TM-align: a protein structure alignment algorithm based on the TM-score. *Nucleic Acids Res,* 33**,** 2302-9.
